# Structural Basis of Glycolytic Control in *Trypanosoma cruzi*: Insights from Enolase and PGI

**DOI:** 10.64898/2026.01.15.699324

**Authors:** Kenneth Austin, Vincent A. Obachi, Florence L. Muzenda, Marakiya T. Moetlediwa, Chelsea Agyei, Timothy Craig, Jan Abendroth, Mary Nguyen, Ngoc Tran, Bart. L. Staker, Tom Edwards, Sandhya Subramanian, Peter. J. Myler, Tawanda Zininga, Krishna K. Govender, Graham Chakafana

**Affiliations:** Chemistry and Biochemistry Department, Hampton University, 200 William R Harvey Way, Hampton, Virginia, 23668, United States; Department of Chemical Sciences, University of Johannesburg, Johannesburg, South Africa; Department of Biochemistry, University of Stellenbosch, Stellenbosch, South Africa; Department of Human Biology, Faculty of Health Sciences, University of Cape Town, Observatory, 7925 Cape Town, South Africa; Seattle Structural Genomics Center for Infectious Disease (SSGCID), Seattle, Washington, USA; UCB BioSciences, Bainbridge Island, WA 98110, USA

**Keywords:** *Trypanosoma cruzi*, Glycolysis, Structure, X-ray crystallography

## Abstract

*Trypanosoma cruzi*, the etiological agent of Chagas disease, depends on glycolysis for ATP production, rendering its glycolytic enzymes attractive targets for therapeutic development. Here, we report the high-resolution crystal structures of two essential glycolytic enzymes, glucose-6-phosphate isomerase (*Tc* PGI, 1.8 Å) and enolase (*Tc* enolase, 2.4 Å) and provide structural and computational analyses to support structure-based drug design. *Tc* PGI adopts a dimeric αβα sandwich fold and features a parasite-specific 53-residue N-terminal extension and a unique C-terminal hook region which both distinguish it from its human ortholog. *Tc* enolase exhibits the conserved (α/β) 8 TIM barrel fold but harbors minor distinct structural deviations, including an extended α17 helix and a structured α1 region, which differentiate it from human isoforms. Both enzymes exhibited high thermal stability, consistent with adaptation to the parasite’s complex life cycle. Structure-based virtual screening using a scaffold with known multi-target potential identified distinct high-affinity inhibitors for each enzyme. Molecular dynamics simulations further confirmed stable enzyme–inhibitor interactions and favorable binding energetics. Collectively, these findings reveal structural signatures unique to *T. cruzi* glycolytic enzymes and lay the groundwork for the development of selective antiparasitic therapeutics.

## Introduction

Chagas disease is a parasitic, systemic, and chronic illness caused by *Trypanosoma cruzi*. According to the World Health Organization (WHO), approximately 70 million people in the Americas live in areas at risk of infection. It is further estimated that 6-7 million individuals are currently infected worldwide, with the majority of cases occurring in Latin America (WHO, 2022). Chagas disease can lead to irreversible damage to the neurological, digestive, and cardiovascular systems, often resulting in long-term disability [1, 2]. The rising prevalence of *T. cruzi* infections underscores the urgent need for novel therapeutic strategies to overcome the limitations of current treatments. Existing drugs, such as nifurtimox (NFX) and benznidazole (BNZ), have reduced efficacy in the chronic phase of the disease and are associated with serious side effects, including neurological and gastrointestinal complications, as well as growing concerns over drug resistance [3, 4]. Despite its significant health burden, Chagas disease remains a neglected tropical disease (NTD), with no approved vaccines and limited therapeutic options. This reality highlights the critical need for intensified research into novel drug targets and the development of more effective treatments that can address all stages of infection.

While protein kinases (PKs) have emerged as promising drug targets in trypanosomatids due to their critical regulatory roles [5, 6], targeting glycolytic enzymes offers a complementary approach to disrupting parasite metabolism. Indeed, glycolytic enzymes are increasingly becoming targets in several diseases including cancer [7, 8]. Recently, glycolytic enzymes have also been explored as potential drug targets in infectious diseases including tuberculosis [9, 10]. During infection, *T. cruzi* relies on glycolysis of host-derived glucose for ATP production. This dependence on glucose metabolism makes glycolytic enzymes attractive targets for therapeutic intervention. In *T. cruzi*, glycolysis is compartmentalized within specialized organelles known as glycosomes, which house the first seven enzymes of the pathway (Figure 1A) [11, 12]. Under aerobic conditions, glycosomal enzymes convert glucose to 3-phosphoglycerate, which is subsequently metabolized to pyruvate in the cytosol, resulting in ATP generation [13, 14]. As such, glycolysis in trypanosomatids is a promising avenue for drug discovery notably because of this compartmentalization of glycolytic enzymes and the essentiality of the pathway to parasite survival [15–17]. Given the parasite’s reliance on this pathway, glycolytic enzymes such as enolase and glucose-6-phosphate isomerase (PGI) present attractive targets for therapeutic intervention. Understanding the structures of these parasitic enzymes is essential for guiding the development of targeted inhibitors against *T. cruzi*, enabling structure-based drug design with greater specificity and efficacy.

**Figure 1.**
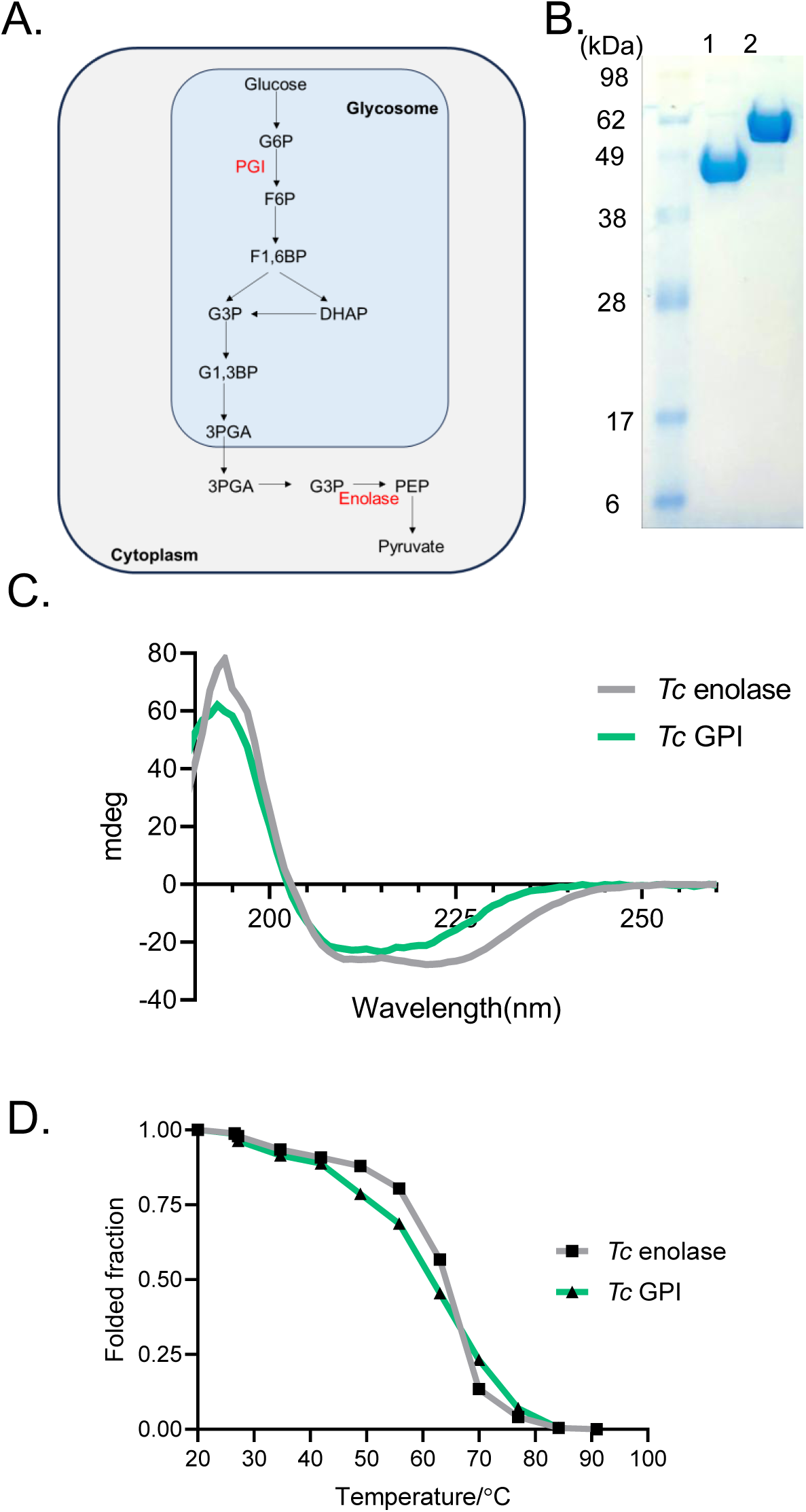
Biochemical characterization of *Tc* PGI and *Tc* enolase. (A) Schematic of the *T. cruzi* glycolytic pathway showing target enzymes PGI and enolase. (B) SDS-PAGE analysis of purified recombinant *Tc* enolase and *Tc* PGI. (C) Far-UV CD spectra and (D) thermal stability analyses of *Tc* PGI and *Tc* enolase.

PGI plays a key role in *T. cruzi* energy metabolism by catalyzing the reversible isomerization of D-glucose-6-phosphate (G6P) to D-fructose-6-phosphate (F6P) [18]. This reaction is essential not only for glycolysis (the parasite’s primary source of ATP) but also for linking glycolysis to the pentose phosphate pathway (PPP). Through this connection, PGI enables the recycling of F6P back into G6P, serving as a substrate for glucose-6-phosphate dehydrogenase, the first step in the PPP [19]. This dual role underscores the enzyme’s metabolic importance and its potential as a therapeutic target. Disrupting PGI activity could impair both energy production and biosynthetic pathways, making it a promising candidate for drug development. A detailed understanding of PGI structure, including how it interacts with G6P substrate is therefore critical for the rational design of selective inhibitors.

Enolase catalyzes the dehydration of 2-phosphoglycerate (2PG) to phosphoenolpyruvate (PEP) in glycolysis and has emerged as a promising therapeutic target due to its additional roles in various cellular processes [20, 21]. Beyond its metabolic function, enolase is considered a moonlighting protein, involved in activities such as mitochondrial tRNA targeting and plasminogen binding [22]. Although these multifunctional roles may add complexity to its targeting, they also broaden its therapeutic potential. Notably, enolase enzymes have already been explored as drug targets in diseases like tuberculosis, where 2-aminothiazole compounds have been shown to inhibit its activity [9]. Previous studies in *Mycobacterium tuberculosis* also demonstrated that altering enolase expression levels modulates drug sensitivity i.e. overexpression confers resistance, while reduced expression enhances susceptibility to inhibition [9]. Additionally, the therapeutic potential of enolase in other pathogens, including *Plasmodium* and *Candida* species has also been demonstrated [23, 24]. Given its essential role in *T. cruzi* metabolism, dysregulating enolase function may impair parasite survival. Furthermore, given that enolase is an immunogenic protein in some infections it may also contribute to host immune recognition [25]. As such, there is need for continued exploration of enolase as a target for novel anti-trypanosomal therapies.

Understanding the three-dimensional structures of glycolytic enzymes is critical for elucidating their functional mechanisms and guiding the design of selective inhibitors. While inhibitors targeting glycolytic enzymes have been identified in other trypanosomatids, such as *T. brucei* and *Leishmania* spp [17, 26], progress in *T. cruzi* has lagged. Nonetheless, studies in related species suggest that even subtle structural differences between mammalian and parasite enzymes can be exploited for selective drug development. Additionally, the unique localization of glycolytic enzymes within glycosomes in *T. cruzi* further enhances their attractiveness as therapeutic targets. In this study, we determined the three-dimensional crystal structures of glucose-6-phosphate isomerase (PGI) and enolase from *T. cruzi*, offering detailed insights into their structural organization. Molecular docking analyses were also conducted to model substrate interactions, and high-throughput virtual screening was used to identify candidate small-molecule inhibitors. These efforts, complemented by molecular dynamics simulations, led to the identification of promising compounds for further investigation. This study deepens the understanding of *T. cruzi* glycolysis from a structural perspective and provides a foundation for structure-based drug discovery aimed at kinetoplastid parasites.

## Materials and methods

### Protein production

The full-length *Tc* enolase (UniProt: Q4DZ98; residues 1-429) and *Tc* PGI (UniProt: O61113; residues 1-608) genes were PCR-amplified using primers listed in Tables S1 and S2, respectively. Both genes were cloned into the pBG1861 vector, which encodes an N-terminal hexahistidine (His₆) tag to facilitate purification. The resulting plasmids were transformed into chemically competent *E. coli* cells. Protein expression was tested, and large-scale production was carried out in 2-liter cultures grown in Terrific Broth at 37°C until mid-log phase, followed by induction with IPTG and incubation at 18°C overnight. Recombinant *Tc* enolase and *Tc* G6PI were purified using a two-step protocol comprising immobilized metal affinity chromatography (IMAC) followed by size-exclusion chromatography (SEC). Frozen bacterial pellets (∼25 g) were thawed and resuspended in 200 mL lysis buffer containing 25 mM HEPES (pH 7.0), 500 mM NaCl, 5% glycerol, 0.5% CHAPS, 30 mM imidazole, 10 mM MgCl₂, 1 mM TCEP, 250 µg/mL AEBSF, and 0.025% sodium azide. Cells were lysed by sonication, followed by treatment with benzonase (20 mL, 25 U/µL) and incubation at room temperature for 45 minutes. The lysate was clarified by centrifugation at 10,000 rpm for 1 hour at 4°C (Sorvall, Thermo Scientific). The clarified supernatant was loaded onto a 5 mL Ni-NTA HisTrap FF column (GE Healthcare) pre-equilibrated with binding buffer (25 mM HEPES, pH 7.0, 500 mM NaCl, 5% glycerol, 30 mM imidazole, 1 mM TCEP, 0.025% sodium azide). After washing with 20 column volumes of the same buffer, proteins were eluted using a linear gradient of imidazole (up to 250 mM) over 7 column volumes. Peak fractions were pooled, concentrated to 5 mL, and further purified by SEC on a Superdex 75 column (GE Healthcare) equilibrated with 20 mM HEPES (pH 7.0), 300 mM NaCl, 5% glycerol, and 1 mM TCEP. Protein elution profiles were monitored by UV absorbance and verified by SDS-PAGE. The peak fractions were collected and analysed using SDS-page for the protein of interest. Purified *Tc* enolase eluted as a single, monodisperse peak at the expected ∼48 kDa molecular weight and >90% purity. Peak fractions were pooled and concentrated to 47.6 mg/mL using an Amicon centrifugal concentrator (Millipore). *Tc* G6PI likewise eluted as a symmetrical, monodisperse peak corresponding to the expected ∼63 kDa species and >90% purity; pooled peak fractions were concentrated to 75.3 mg/mL (Amicon, Millipore). Both proteins were aliquoted (110 µL), flash-frozen in liquid nitrogen, and stored at −80 °C until use.

### Circular dichroism spectroscopy and thermal stability analysis

The secondary structure content and thermal stability of *Tc* enolase and PGI enzymes were analyzed using far-UV circular dichroism (CD) spectroscopy on a JASCO CD spectropolarimeter (JASCO International, Japan). Far-UV CD spectra were recorded to assess overall secondary structure using a scan speed of 100 nm min⁻¹, a 1-nm bandwidth, and seven accumulations, with buffer spectra subtracted as baseline controls. Thermal denaturation experiments were also performed to evaluate the stability of protein secondary structure. CD signals were monitored at 222 nm, a wavelength sensitive to α-helical content, while the temperature was increased monotonically from 20 to 90 °C at a rate of 0.5 °C min⁻¹. Ellipticity measurements were recorded at 7 °C intervals, with an equilibration period of 60 s at each temperature to ensure thermal equilibrium prior to data acquisition.

The fraction of folded protein at any given temperature was calculated according to the equation [27]:

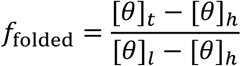

where [*θ*]_*t*_represents the molar ellipticity at a given temperature, [*θ*]_*h*_is the molar ellipticity at the highest temperature (fully unfolded state), and [*θ*]_*l*_is the molar ellipticity at the lowest temperature (fully folded state). The apparent melting temperature (T_m_) was determined as the midpoint of the unfolding transition by fitting the temperature-dependent ellipticity data to a two-state unfolding model using GraphPad Prism software.

### Crystallization

*Tc* enolase and *Tc* G6PI crystals were screened in 96-well plates using commercial crystal screens, JB Screen JCSG++ HTS (Jena Bioscience) and MCSG1 (Molecular Dimensions) crystal screens. Equal volumes of protein solution (0.4 µl) and precipitant solution were set up at 290 K against reservoir (80 µl) in sitting-drop vapor-diffusion format. The crystals were harvested and cryoprotected with 20% ethylene glycol prior to data collection (Table S3).

#### Data collection and processing

Data were collected at 100 K on beamlines 21-ID-FG and 5.0.3 at the Advanced Photon Source, Argonne National Laboratory (Table S4). Data were integrated with XDS and reduced with XSCALE [28]. Raw X-ray diffraction images for 4QFH and 4G7F are available at the Integrated Resource for Reproducibility in Macromolecular Crystallography at https://www.proteindiffraction.org.

### Structure solution and refinement

The structures of *Tc* enolase and *Tc* PGI were determined by molecular replacement with Phaser [29] from the CCP4 suite of programs [30] using domains of PDB entries of 1OEP and 2O2C, respectively as search models. The quality of each structure was checked using MolProbity. Data-reduction and refinement statistics are shown in Table 1. Coordinate and structure factors have been deposited into the Protein Data Bank (PDB) as entries 4G7F (*Tc* enolase) and 4QFH (*Tc* PGI).

**Table 1.** Structure solution and refinement.

|  | *Tc* enolase | *Tc* PGI |
| --- | --- | --- |
| Resolution range (Å) | 19.809–2.400 (2.5263–2.4000) | 29.191–1.800 (1.8205–1.8000) |
| Completeness (%) | 100.0 | 98.7 |
| $\sigma$ cutoff | $F > 1.36\sigma(F)$ | $F > 1.36\sigma(F)$ |
| No. of reflections, working set | 19774 (2663) | 108307 (3429) |
| No. of reflections, test set | 993 (143) | 5596 (180) |
| Final $R_{\text{cryst}}$ | 0.189 (0.2181) | 0.139 (0.2345) |
| Final $R_{\text{free}}$ | 0.250 (0.3025) | 0.175 (0.2978) |
| No. of non-H atoms |  |  |
| Protein | 2997 | 9548 |
| Ion | 1 |  |
| Ligand | 4 | 32 |
| Water | 89 | 1136 |
| Total | 3091 | 10716 |
| R.m.s. deviations |  |  |
| Bonds (Å) | 0.014 | 0.007 |
| Angles (°) | 1.467 | 1.012 |
| Average $B$ factors (Å<sup>2</sup>) | | |
| Protein | 40.3 | 20.9 |
| Ion | 30.8 |  |
| Ligand | 26.8 | 22.4 |
| Water | 34.3 | 31.0 |
| Ramachandran plot |  |  |
| Most favoured (%) | 98 | 98 |
| Allowed (%) | 1 | 1 |
| Outlier (%) | 1 | 1 |

### High throughput virtual screens

The crystal structures of *Tc* enolase (PDB ID: 4G7F) and *Tc* PGI (PDB ID: 4QFH) were retrieved from the Protein Data Bank. Potential binding sites were identified using Schrödinger’s SiteMap module, which assigns druggability scores based on site size, hydrophobicity, and enclosure [31]. To validate the SiteMap results, blind docking was performed using PyRx AutoDock, encompassing the entire protein structure. For *Tc* enolase, the blind docking grid was centered at x=-24.69, y=-21.84, z=-19.84 with dimensions x=67.52, y=83.81, z=66.34. For *Tc* PGI, the grid was centered at x=24.05, y=41.59, z=12.93 with dimensions x=84.29, y=97.76, z=75.05. The blind docking confirmed that the highest-scoring SiteMap site was the primary binding pocket, as all potential inhibitors bound to the same site, validating the SiteMap predictions. An alternative site with lower SiteMap scores yielded inferior docking scores and was excluded. Enzyme structures were prepared using the Chimera AutoDock tool, where they were minimized with Gasteiger charges to optimize geometry. Additional processing in PyRx converted the structures into AutoDock-compatible macromolecules. A set of 21 ligands, including 2-PGA and compounds from studies on enolase and PGI inhibitors [32], were prepared in Chimera AutoDock by adding hydrogens and charges, then saved as mol2 files. In PyRx, ligands were minimized using the Universal Force Field (UFF) with conjugate gradients over 200 steps and converted to AutoDock ligand format (pdbqt). Molecular docking was performed using AutoDock Vina within PyRx, targeting the validated binding sites. Key amino acid residues within 5 Å of the binding sites were identified using Chimera. For *Tc* enolase, these included Arg15, Gly38, Ala39, Lys155, His156, Gln164, Asp207, Glu208, Asp243, Ser247, Glu248, Glu291, Asp292, Asp318, Asp319, Lys343, Gln346, Arg372, and Ser373. For *Tc* PGI (4QFH), residues included Arg145, Ile206, Gly207, Gly208, Ser209, Ala258, Ser259, Lys260, Thr261, Phe262, Thr264, Thr267, Phe320, Gly325, Gly326, Arg327, Tyr328, Ser329, Ile334, Gln408, Glu412, Gln565, Val568, and Lys572.

### Molecular Dynamics Simulations and Post-MD

Molecular dynamics (MD) simulations were conducted using AMBER 20 with the pmemd.cuda module to enable GPU acceleration. Protein-ligand complexes derived from the top-scoring docking poses were prepared using the ff14SB force field for proteins (*Tc* enolase and PGI) and GAFF for ligands, with ligand partial charges assigned via AM1-BCC using antechamber. Each system was solvated in a truncated octahedral TIP3P water box with a 10 Å buffer, and neutralized with Na⁺ or Cl⁻ ions to achieve a physiological salt concentration of 0.15 M. Initial energy minimization was performed in two stages: 5,000 steps of steepest descent followed by 5,000 steps of conjugate gradient minimization, terminating when the system reached a force threshold of 10 kcal/mol/Å². Equilibration was carried out under NVT conditions for 500 ps at 300 K using a Langevin thermostat (collision frequency of 2.0 ps⁻¹), followed by NPT equilibration for 1 ns at 1 bar using a Monte Carlo barostat. Production MD simulations were run for 120 ns with a 2 fs integration time step, employing SHAKE to constrain bonds involving hydrogen atoms. Periodic boundary conditions were applied, with long-range electrostatics treated using the particle mesh Ewald (PME) method and a 10 Å non-bonded cutoff. Trajectories were saved every 10 ps. Post-simulation analysis was performed using cpptraj to calculate root-mean-square deviation (RMSD) and root-mean-square fluctuation (RMSF), providing insights into the structural stability and residue-level flexibility of the protein-ligand complexes. Binding free energy calculations were performed using the MM/GBSA method implemented in MMPBSA.py, analyzing 120 ns trajectories sampled every 10 frames. Binding free energy (ΔG_bind) was decomposed into van der Waals (ΔE_vdW), electrostatic (ΔE_elec), and solvation energy (ΔG_solv) components. Solvation energy was computed using the Generalized Born model (igb=2) with a salt concentration of 0.15 M and SASA-based nonpolar contributions (1.4 Å probe radius; 0.0072 kcal/mol/Å² surface tension). Entropic contributions were not included due to computational cost. A total of 100 snapshots at 100 ps intervals were used for the MM/GBSA calculations.

## Results

### Biochemical characterization of *Tc* PGI and *Tc* enolase

Recombinant *Tc* PGI and *Tc* enolase were successfully expressed in *E. coli* and purified using a two-step protocol involving immobilized metal affinity chromatography followed by size-exclusion chromatography. SDS-PAGE analysis confirmed the purity and expected molecular weights of the proteins, with *Tc* PGI migrating at approximately 63 kDa and *Tc* enolase at 48 kDa (Figure 1B). To evaluate secondary structure content and folding quality, far-UV circular dichroism (CD) spectroscopy was performed on both enzymes. The CD spectrum of *Tc* PGI exhibited characteristic double minima near 208 nm and 222 nm, consistent with a predominantly α-helical secondary structure (Figure 1C). *Tc* enolase displayed a CD spectrum indicative of a mixed α/β structure, aligning with its known (α/β) 8-barrel fold (Figure 1C). These spectra confirm proper folding of both proteins following purification. Thermal denaturation profiles, monitored by changes in ellipticity at 222 nm as a function of temperature, provided insights into protein stability. Both proteins displayed high stability under heat, as *Tc* PGI exhibited a melting temperature (Tm) of approximately 65°C, while *Tc* enolase had a Tm of ∼58°C (Figure 1D). These findings indicate that *T. cruzi* glycolytic enzymes display high thermal stability

### Crystal structure of *Tc* enolase

The crystal structure of *Tc* enolase reveals a bilobal architecture that is highly conserved among members of the enolase superfamily (Figure 2A). The enzyme is composed of two structurally distinct domains: a smaller N-terminal domain (primary domain) and a larger C-terminal domain (secondary domain). The primary domain consists of three β-strands (β1-β3) arranged as a mixed β-sheet flanked by five α-helices (α1-α5). This domain forms a core component of the domain interface and serves as a structural scaffold that supports the positioning of the larger secondary domain. In contrast, the secondary domain adopts a canonical (α/β) 8 TIM barrel fold, comprising eight parallel β-strands surrounded by eight α-helices, which form the central catalytic core of the enzyme. The active site is located at the top of the TIM barrel, at the interface between the two domains within a single subunit, and contains a catalytically essential Mg²⁺ ion coordinated by conserved residues (Figure 2B). The topology diagram (Figure 2B) outlines the sequential arrangement of helices (H1–H17) and β-strands (B1–B12), highlighting the asymmetric distribution of secondary structure elements between the two domains. The α5 and α6 helices from the N-terminal domain are positioned near the active site and may play a role in regulating substrate access or stabilizing the catalytic conformation. *Tc* enolase catalytic site features the characteristic octahedral geometry, with Mg²⁺ coordinated by D^243^, E^291^, and D^318^, residues critical for stabilizing the enolate intermediate during catalysis (Figure 2D). Additional catalytic residues, including H^159^ and K^341^, are well-positioned to support proton abstraction and substrate binding. Importantly, substrate binding and catalysis are mediated by residues contributed from both domains, consistent with the enolase function of converting 2-phosphoglycerate (2-PGA) to phosphoenolpyruvate (PEP).

**Figure 2.**
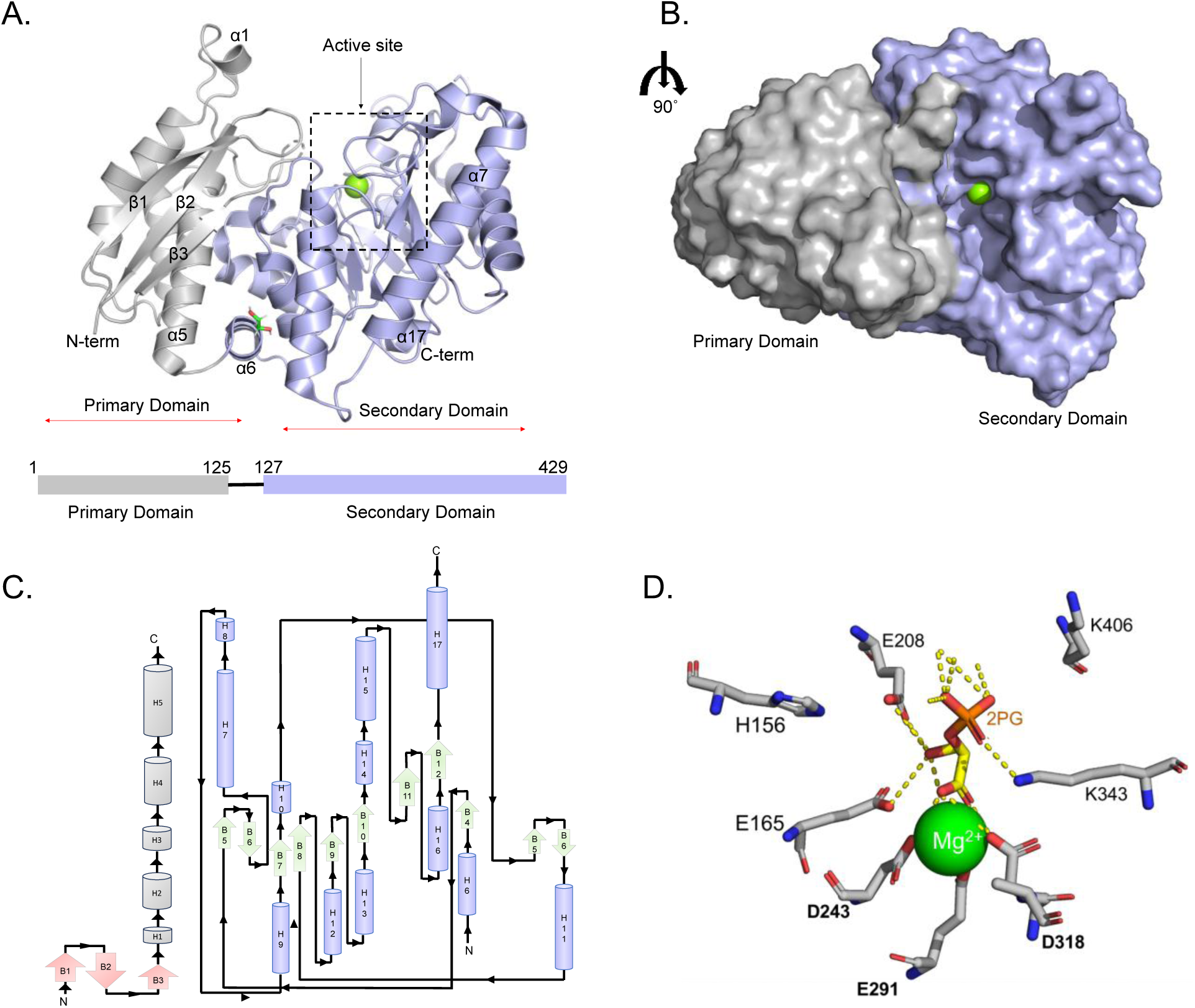
Structural and topological characterization of *T. cruzi* enolase. (A) Ribbon diagram of the crystal structure of *Tc* enolase (PDB ID: 4G7F) and the domain organization of *Tc* enolase. (B) Surface representation of Tc enolase showing domain interface. (C) Topology diagram of *Tc* enolase showing the organization and connectivity of secondary structure elements. (D) *Tc* enolase active site. Metal binding sites are in bold.

### Comparative structural analysis of *Tc* enolase

To identify species-specific structural features of *Tc* enolase, we performed both MSA and three-dimensional structure comparisons with other eukaryotic and bacteria enolases (Figure 3, Figure S1-4). Structural alignments revealed a high degree of conservation across the enolase family, particularly in the core (α/β) 8-barrel catalytic domain and in in active site residues. However, distinct differences were observed in the peripheral regions of the enzyme. Structural superposition of *Tc* enolase with human enolase isoforms (Figure 3A) confirmed the preservation of the classical enolase fold, including the α/β-barrel architecture and active site topology. Notably, the residues that stabilize Mg^2+^ binding (P^263^, D^264^, and K^261^) are conserved across species. Nonetheless, a notable structural feature of *Tc* enolase is the extended C-terminal α-helix (α17), which is less prominent in human isoforms. Structural divergence was also evident in peripheral loop regions and surface charge distributions, which may underlie differences in protein-protein interactions or regulation by host-derived factors. The α1 region in *Tc* enolase also forms a defined α-helical motif, whereas the corresponding region in human enolase isoforms adopts a loop conformation (Figure 3A). This parasite-specific secondary structure element may contribute to differences in protein stability, surface topology, or interaction with parasite-specific partners, and thus represents a unique structural feature that could be exploited for selective drug targeting.

**Figure 3.**
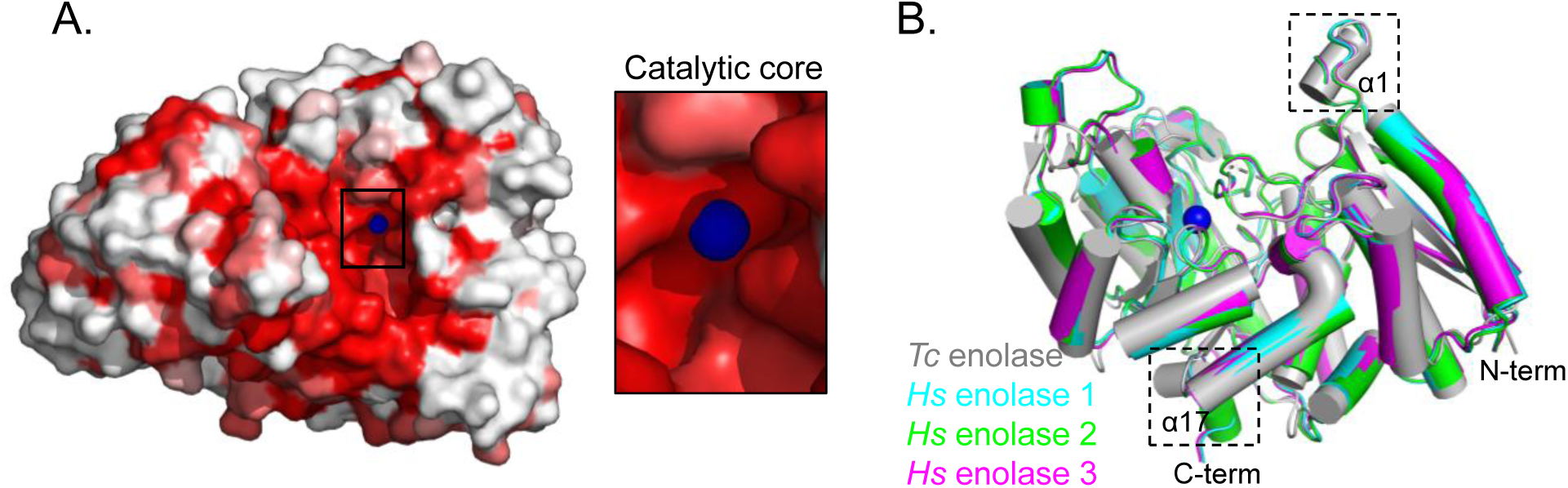
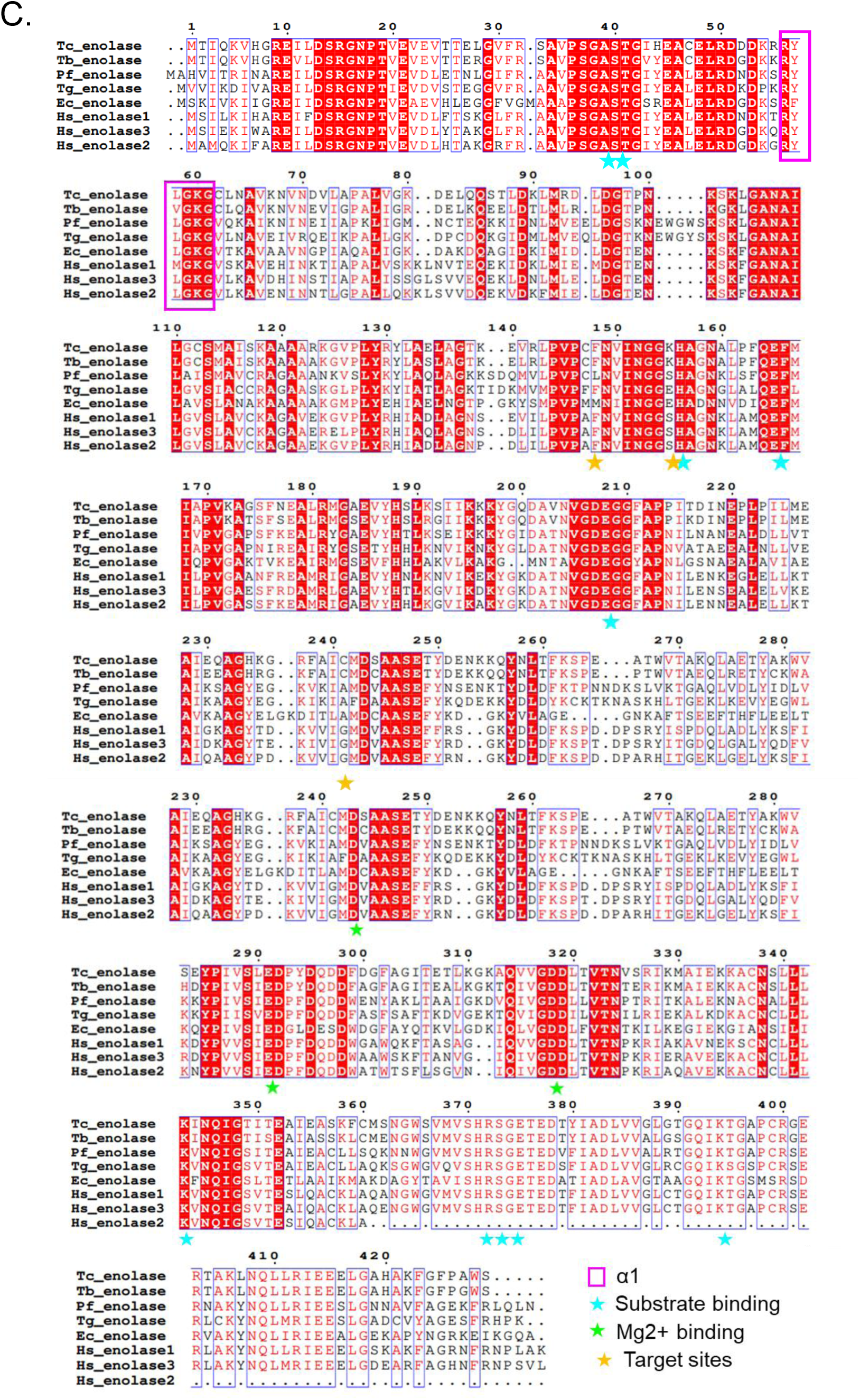
Structural comparison of *Tc* enolase with human enolase isoforms. (A) Mapping of structural divergence between *Tc* enolase and 62 enolase structures in the PDB (accession numbers in Figure S1-4). Regions with higher structural deviation are colored red, while conserved regions are shown in white. (B) Structural superposition of *Tc* enolase with human enolase isoforms 1, 2 and 3. (C) Multiple sequence alignment of select enolases from T.cruzi (Tc enolase), T. brucei (Tb enolase), P. falciparum (Pf enolase), T. gondii (Tg enolase), E. coli (Ec enolase) and Human (Hs enolase 1-3).

To further assess structural variability, we mapped divergence between *Tc* enolase and 62 structurally resolved enolases from the Protein Data Bank (Figure 3B). This analysis showed that while the catalytic core remains highly conserved, the greatest variation is localized to peripheral loops and helices. The high conservation of the catalytic core supports functional equivalence of *Tc* enolase with other members of the enolase family. In contrast, subtle differences in side-chain orientation and active-site cavity accessibility may provide opportunities for selective inhibition of the parasite enzyme. Whether and how these peripheral structural variations contribute to parasite-specific function or druggability warrants further investigation. Multiple sequence alignment (MSA) with representative eukaryotic and bacterial enolases (Figure 3C; Figures S1-S5) also revealed a high degree of sequence conservation across the enolase family, particularly across residues involved in substrate binding and metal coordination (Figure 3C). Interestingly, *Tc enolase* retains a set of chemically reactive residues, Lys155, Cys147, and Cys241, that have previously been implicated in the design of selective Trypanosoma brucei enolase inhibitors. The presence of these residues in Tc enolase, combined with their local structural environment, further supports the feasibility of targeting conserved catalytic machinery through parasite-specific inhibitory strategies. [33].

### Crystal structure of *Tc* PGI

Glucose-6-phosphate isomerase (PGI) from *T. cruzi* exists predominantly as a homodimer in the cytoplasm, although monomeric forms have been isolated under extracellular conditions [34]. The dimer, with an approximate molecular weight of 126 kDa, comprises two identical subunits arranged in a head-to-tail fashion (Figure 4A, B). Dimerization is mediated by extensive interactions, particularly through interlocking protrusions on each subunit that form a distinctive “hugging” interface. The active site of each protomer is positioned near this interface, nestled between the N- and C-terminal domains, highlighting the functional importance of oligomerization (Figure 4C). Each monomer consists of two domains connected by a flexible hinge: a larger N-terminal domain and a smaller C-terminal domain, both adopting characteristic αβα topology (Figure 4C, D). The N-terminal domain contains six mixed β-strands flanked by α-helices, contributing to the catalytic core, while the C-terminal domain includes seven β-strands and houses structural elements critical for inter-subunit communication (Figure 4). *Tc* PGI contains a C-terminal peroxisomal targeting signal type 1 (PTS1), composed of the tripeptide Ser-His-Leu (SHL) (Figure 5D, S8). The PTS1 signal mediates localization of the enzyme to the glycosome and distinguishes it from its cytosolic homolog [16]. A hallmark feature of *Tc* PGI is the extended C-terminal “hook” region, which wraps around the adjacent subunit and likely contributes to dimer stability and potential allosteric regulation.

**Figure 4.**
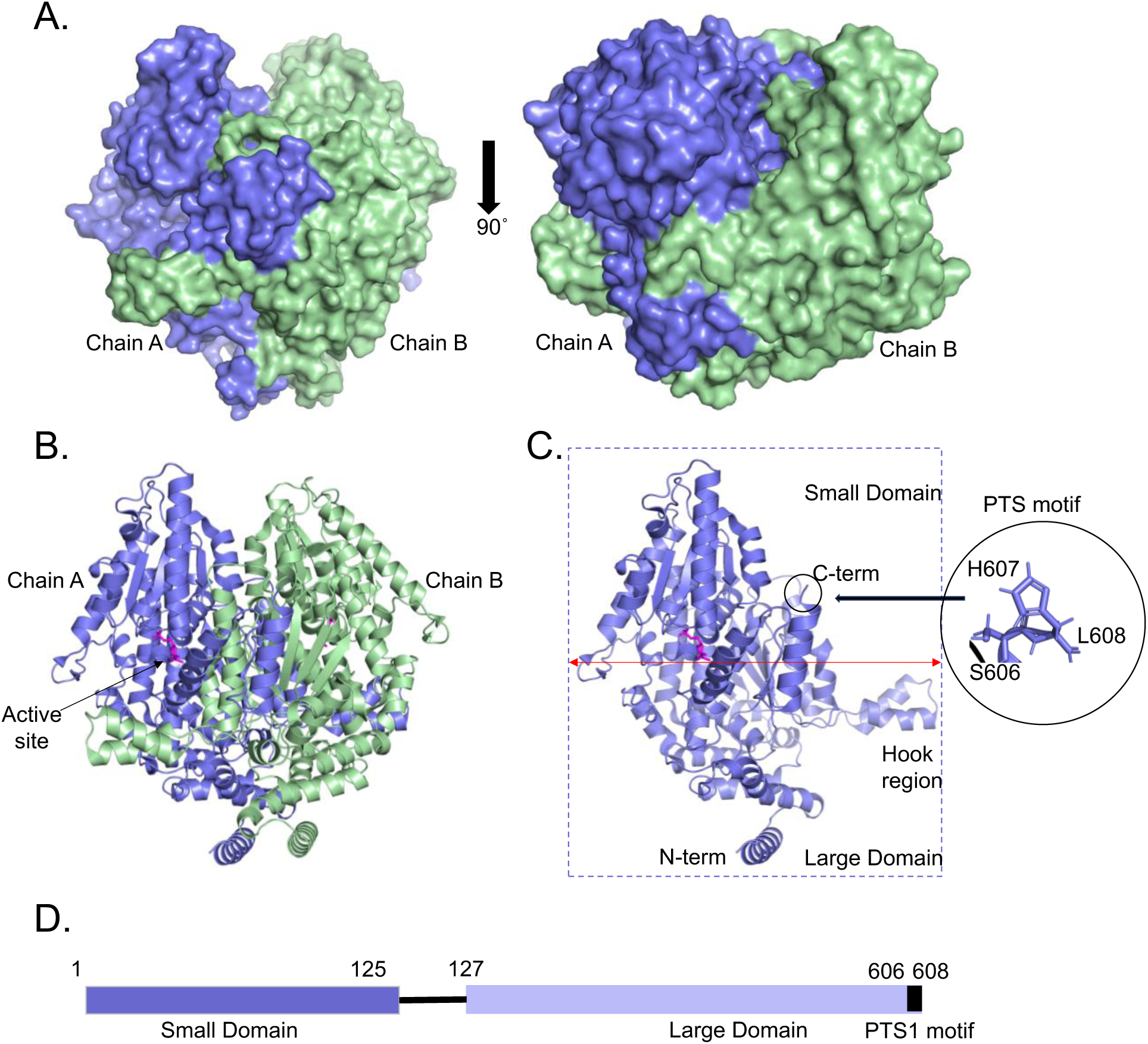
Structural architecture of *Tc* PGI. (A) Surface representation of Tc PGI homodimer, composed of Chain A (blue) and Chain B (green), shown in two orientations to highlight the dimer interface. (B) Ribbon representation of the *Tc* PGI dimer, colored by chain. (C) Structural features of a single *Tc* PGI monomer. (D) Domain organization of a single *Tc* PGI monomer.

**Figure 5.**
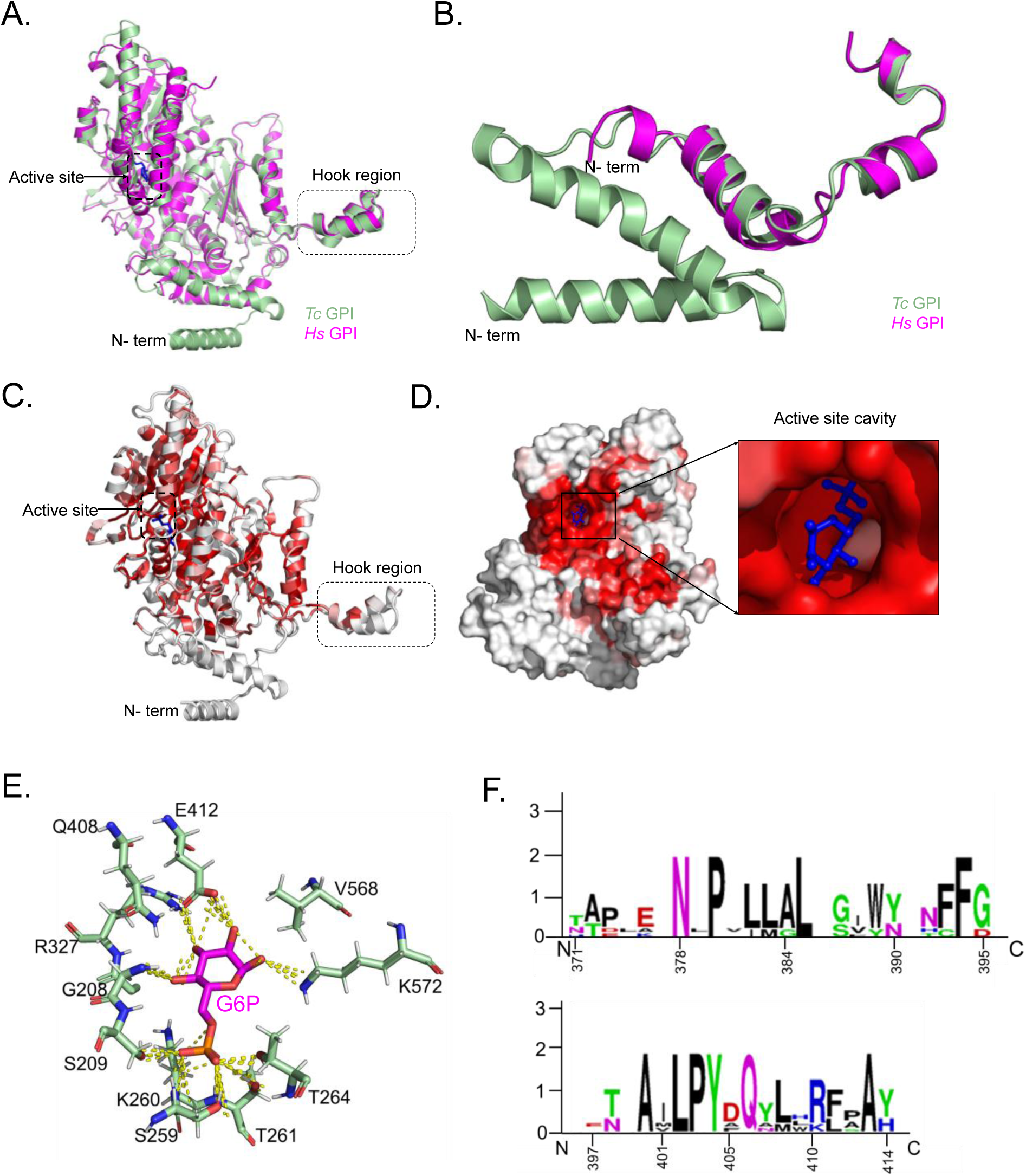
Comparative structural analysis of *Tc* PGI with other PGIs. (A) Structural superposition of *Tc* PGI and human PGI. (B) Structural divergence of *Tc* PGI compared to other PGI structures in PDB. Highly divergent regions are shown in red, while conserved regions are in white. (C) Close-up view of the *Tc* PGI active site highlighting key residues involved in substrate binding and catalysis. (D) Comparative analyses of Hook residues from Tc PGI and other eukaryotic and bacterial PGIs.

### Structural comparison of *Tc* PGI with other PGIs

To elucidate species-specific structural features of *Tc* PGI, we conducted comparative structural analyses with PGI homologs from other organisms. Structural superposition with human PGI (*Hs* PGI) revealed a high degree of conservation in the core αβα sandwich fold, a hallmark of eukaryotic PGIs (Figure 5A). Notably, *Tc* PGI possesses a hydrophobic 53-residue N-terminal extension that is absent in *Hs* PGI (Figure 5A, B, S9). A similar extension is also present in *T. brucei* PGI (Figure S9), suggesting that this feature is conserved among trypanosomatid PGIs. While the functional role of this parasite-specific N-terminal extension remains to be defined, its absence in the human homolog suggests a potential role in parasite-specific regulation. Additionally, while the catalytic core remains structurally conserved, loop regions adjacent to key residues such as Glu314 and Arg327 display conformational variability, suggesting possible differences in substrate recognition or catalytic dynamics.

Further analysis using ENDSCRIPT to compare *Tc* PGI with 28 other PGI structures in the Protein Data Bank (PDB) (Figure 5C-D, Figure S6-9) revealed significant divergence in the N-terminal region of the large domain and within the C-terminal hook region. Nonetheless, there was high conservation within the active site across various species (Figure 5D). We further explored the binding mode of glucose-6-phosphate (G6P) into the *Tc* PGI active site. Key residues identified within 4 Å of the ligand included Gly208, Ser209, Ser259, Thr261, Thr264, Arg327, Glu 412 and Lys572 which formed hydrogen bonds with G6P substrate (Figure 5E). Additional hydrophobic interactions were observed with Lys260. Mechanistic interpretation of the binding mode suggests that Arg327 may act analogously to a lysine or histidine in catalyzing ring opening by donating a proton to the ring oxygen, thereby stabilizing the open-chain form of the sugar. Subsequently, Glu412 likely serves as a catalytic base, abstracting the proton from carbon 2 to generate a cis-enediolate intermediate (Figure 5E). The same Glu412 residue may then act as a proton donor to carbon 1, while Arg327 retrieves the proton from the ring oxygen to promote ring closure. This two-residue mechanism is consistent with previously proposed models for PGI catalysis and is supported by the proximity and polarity of these residues observed in the docking analysis. The hook region (comprising residues 371-414) demonstrated a degree of variation. Comparative sequence analyses across protozoan parasites (*Toxoplasma gondii*, *Plasmodium falciparum*, *Trypanosoma brucei*), bacteria (*Escherichia coli*), and human PGI isoforms revealed notable divergence within this region. Despite this variability, several residues within the core of the hook, most notably N378, P379, and a hydrophobic cluster spanning L381-L384, were conserved (Figure 5F, S10). These conserved residues are likely critical for stabilizing the dimer interface and maintaining the structural integrity of the enzyme. Importantly, the flanking regions surrounding this conserved core differ between parasite and human PGI which could result in distinct local conformational dynamics at the dimer interface. Such differences may also influence allosteric communication between subunits or modulate access to adjacent cavities, thereby mediating parasite-specific protein-protein interactions within the glycosome.

### *In silico* drug screening

The molecular docking results, summarized in Table 4, highlight the binding affinities of 23 ligands, including the substrates, 2-phosphoglyceric acid (2-PGA) for *Tc* enolase and glucose-6-phosphate for *Tc* PGI. Among the *Tc* enolase ligands, Cpd8 showed the strongest binding affinity at −8.0 kcal/mol, followed by Cpd12 (−7.9 kcal/mol) and Cpd7 (−7.7 kcal/mol). For *Tc* PGI, Cpd14 exhibited the highest affinity, with Cpd21 and Cpd15 ranking next (Figure S10). In comparison, the substrates displayed markedly lower affinities for both enzymes, confirming that the top-performing compounds significantly outperform the substrates.

### Protein-Ligand Interaction Analysis

We then assessed the binding modes for the protein-ligand interactions between the compounds and *Tc* enolase and *Tc* PGI enzymes (Figure 6). We noted that in *Tc* enolase, Cpd8 forms hydrogen bonds with Asp207 and Glu208, and hydrophobic interactions with Arg372, stabilizing its high-affinity binding. Similarly, Cpd12 employs a similar mechanism through which it engages with the same residues, with additional hydrophobic contacts at Lys343 and Ser373, contributing to its −7.9 kcal/mol affinity. Cpd7 interacts with Asp207, Arg372, and Gln346, supporting its strong binding at −7.7 kcal/mol. These interactions align with *Tc* enolase’s active site architecture which is critical for its role in glycolysis. In *Tc* PGI, Cpd15 forms hydrogen bonds with Thr264 and Ile206, complemented by hydrophobic interactions with Lys572 and Trp566. Additionally, Cpd21 interacts with *Tc* PGI’s Lys260 and Phe320 residues, while Cpd14 associates with residues Arg327 and Ser329, contributing to their respective affinities of −7.7 and - 7.6 kcal/mol. These interactions highlight the structural compatibility of these ligands with the enzyme’s active site.

**Figure 6.**
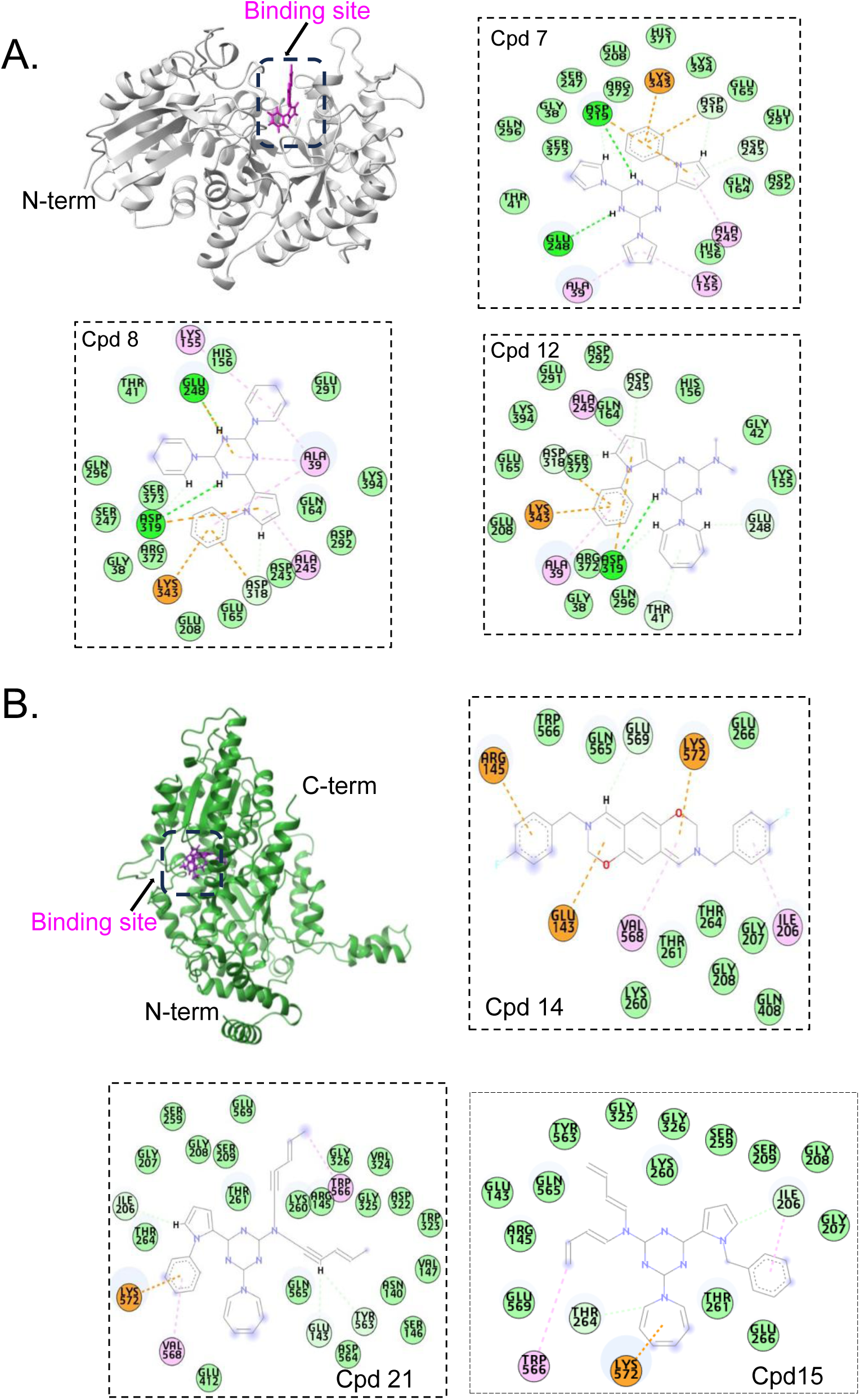
Superimposed inhibitor structures in the active sites of (A) *Tc* enolase complexes with Cpd7, Cpd8, and Cpd12, and (B) *Tc* PGI with Cpd14, Cpd21 and Cpd15, highlighting their binding modes.

### Molecular Dynamics Simulations and Post-MD Analysis

To assess the dynamic stability of the lead compounds within the active sites of *Tc* enolase and PGI, MD simulations were performed for both *apo* forms and selected protein-ligand complexes. For *Tc* enolase, the top three docked ligands (Cpd7, Cpd8, and Cpd12) were simulated alongside the *apo* enzyme. In the case of *Tc* PGI, compounds Cpd14, Cpd15, and Cpd21 were selected for evaluation. The *Tc* enolase RMSD profiles (Figure 7A) revealed that Cpd8 maintained the most stable complex, with an average RMSD of 1.36 Å, outperforming the *apo* form (1.83 Å), Cpd7 (1.55 Å), and Cpd12 (1.42 Å). Despite Cpd12’s competitive docking score, its higher RMSD indicates slightly reduced conformational stability. RMSF analysis (Figure 7B) demonstrated that Cpd8 and Cpd12 induced minimal fluctuations in key catalytic residues-Asp319, Glu248, and Lys343- at 0.80 Å and 0.76 Å, respectively, compared to 0.75 Å for the *apo* form and 0.83 Å for Cpd7. The relatively higher flexibility observed with Cpd7 may compromise its binding consistency. For *Tc* PGI, RMSD results (Figure 7C) showed the *apo* enzyme with the lowest deviation (4.00 Å), closely followed by Cpd14 (4.06 Å). Cpd21 showed moderate deviations (4.50 Å), while Cpd15 had the highest RMSD (4.94 Å), exceeding the generally accepted threshold of 3 Å for stable complexes. This elevated RMSD suggests possible ligand-induced structural rearrangements or suboptimal binding interactions, potentially due to steric hindrance or poor hydrogen bonding. RMSF profiles indicated that Cpd14 stabilized key residues (Glu143, Tyr563, and Ile206) with the lowest fluctuations (1.41 Å), in line with its strong docking score. By contrast, Cpd15 (1.59 Å) and Cpd21 (1.98 Å) induced greater flexibility, suggesting less favorable binding stability compared to the apo form (1.60 Å).

**Figure 7.**
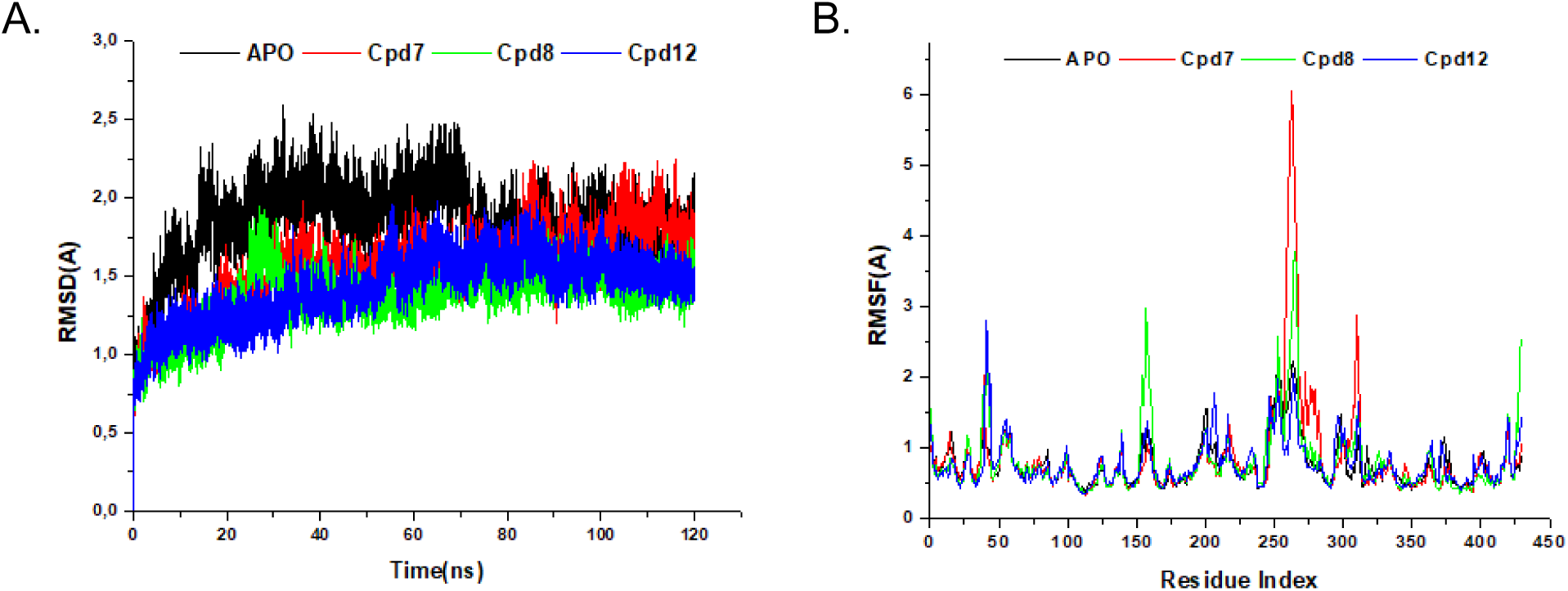
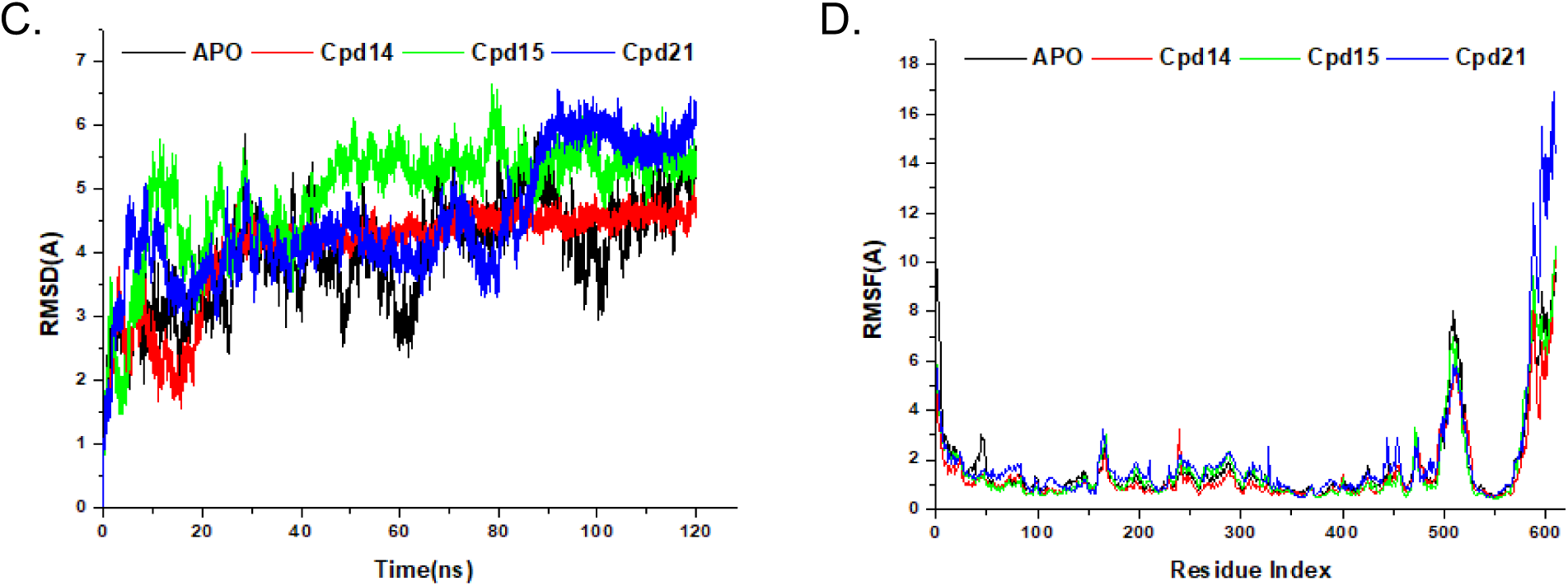
RMSD and RMSF plots for (A, B) *Tc* enolase complexes with apo, Cpd7, Cpd8, and Cpd12, and (C, D) *Tc* PGI complexes with Cpd14, Cpd15, and Cpd21, illustrating structural stability and residue flexibility.

### Binding Free Energy Analysis

Binding free energy analysis using the MM/GBSA method further elucidated the energetic profiles of these complexes, as presented in Table 2. For *Tc* enolase, Cpd8 exhibited the most favourable binding free energy at −42.26 kcal/mol, driven by a significant electrostatic contribution (ΔEelec), despite a high solvation penalty (ΔGsolv). Cpd7 followed with a binding free energy of −39.84 kcal/mol, benefiting from a strong van der Waals contribution (ΔEvdW), while Cpd12 showed the least favorable energy at −11.27 kcal/mol, with weaker contributions across all terms. These results align with the docking and MD stability trends, where Cpd8’s high affinity and stability correlate with its superior binding energy, making it the top candidate for *Tc* enolase.

**Table 2.** Binding free energy profiles (in kcal/mol) for *Tc* enolase (grey) and PGI (green) complexes with best compounds.

| Complex | $\Delta E_{vdW}$ | $\Delta E_{elec}$ | $\Delta G_{gas}$ | $\Delta G_{solv}$ | $\Delta G_{bind}$ |
| --- | --- | --- | --- | --- | --- |
| Cpd7 | -48.20±3.48 | -214.74±12.05 | -262.94±13.30 | 223.11±11.64 | -39.84±5.44 |
| Cpd8 | -27.65±3.48 | -726.44±69.16 | -754.09±74.53 | 712.84±68.15 | -42.26±8.53 |
| Cpd12 | -16.78±12.96 | -118.56±57.65 | -135.35±68.67 | 124.08±60.54 | -11.27±10.42 |
| Cpd14 | -56.74±3.24 | -68.23±13.64 | -124.97±13.65 | 79.70±13.49 | -45.27±4.63 |
| Cpd15 | -37.20±7.92 | -53.90±18.32 | -91.10±17.81 | 63.58±17.64 | -27.52±7.29 |
| Cpd21 | -37.93±8.13 | -92.91±14.87 | -130.85±17.94 | 97.15±15.28 | -33.69±9.88 |

For *Tc* PGI, Cpd14 exhibited the most favorable binding free energy at −45.27 kcal/mol, driven by strong electrostatic and van der Waals interactions, followed by Cpd21 (−33.69 kcal/mol) and Cpd15 (−27.52 kcal/mol). The combination of Cpd14’s superior binding energy and relatively stable RMSD value (4.06 Å) supports its candidacy as the most promising inhibitor, despite the overall high RMSD values observed across all PGI-ligand complexes indicative of potential conformational flexibility or binding-site adaptation. In contrast, Cpd15 showed both the weakest binding energy and highest RMSD (4.94 Å), suggesting reduced stability, while Cpd21 displayed intermediate behavior in both metrics (−33.69 kcal/mol; 4.50 Å).

## Discussion

*T. cruzi*, the etiological agent of Chagas disease, remains a major parasitic health threat in Latin America. Despite its clinical relevance, existing treatment options are limited in efficacy and associated with significant side effects, underscoring the urgent need for novel therapeutic strategies[3, 4]. This study provides structural insights into two essential glycolytic enzymes from *T. cruzi;* glucose-6-phosphate isomerase and enolase. These enzymes are integral to the parasite’s ATP-generating glycolytic pathway and therefore represent promising candidates for antiparasitic drug development[11]. Our initial analyses evaluated the thermal stability of the secondary structures of *Tc* PGI and enolase using thermal denaturation assays (Figure 1). Both enzymes demonstrated high thermal stability, with melting temperatures of 65°C for *Tc* PGI and 58°C for *Tc* enolase (Figure 1). These findings are consistent with the biology of *T. cruzi*, which cycles between insect vector and mammalian host environments that impose varying thermal and oxidative stress[35, 36]. The parasite’s ability to survive in these fluctuating conditions is underpinned by a proteome enriched in thermostable and stress-resilient proteins. Indeed, thermal resilience is a common feature among kinetoplastid proteins, including chaperones, glycolytic enzymes, and redox regulators from *Trypanosoma brucei, Plasmodium* and *Leishmania* species, which similarly exhibit elevated stability to support parasite survival under hostile conditions[37–40].

The crystal structure of *Tc* enolase reveals a bilobal configuration characteristic of the enolase superfamily, with a conserved catalytic core embedded within a (α/β)_8 TIM barrel fold (Figure 2). The primary domain, comprising a compact β-sheet flanked by α-helices, forms a stabilizing interface that supports dimerization and positions the secondary domain, which houses the active site. This domain arrangement reflects the evolutionary conservation of structure-function relationships across enolase orthologs and underscores the enzyme’s central role of catalyzing the dehydration of 2-PGA to PEP in glycolysis. Comparative structural analyses with 62 enolase homologs further confirm that the catalytic core is highly conserved, as expected given the essential function of enolase across diverse organisms (Figure 3). Residues involved in Mg²⁺ coordination (D^264^, E^295^, D^320^) and proton abstraction (H^159^, K^341^) are well-conserved, maintaining the classical octahedral geometry critical for enolate intermediate stabilization during catalysis. However, differences in surface electrostatics, side-chain orientations near the active site, and the openness of the catalytic pocket may affect substrate access or inhibitor binding, offering a rationale for structure-based design of species-specific inhibitors [41]. Despite its conserved fold, *Tc* enolase exhibits an extended α17 helix at the C-terminus and the distinct α-helical conformation in the α1 region are absent or less defined in human isoforms (Figure 3). Further studies are required to determine whether these differences provide potential footholds for the development of selective inhibitors that spare the host enzymes.

The crystal structure of *Tc* PGI was resolved at 1.8 Å resolution, revealing a dimeric αβα sandwich fold typical of eukaryotic PGIs (Figure 4). Notably, *Tc* PGI contains a 53-residue N-terminal extension and a unique C-terminal hook region which distinguishes it from its human homolog (Figure 5; [42]). Whether these parasite-specific elements present valuable opportunities for the development of selective inhibitors warrants further investigation. The hook region is likely to be conformationally flexible, potentially hindering the formation of stable ligand-binding pockets. Similarly, the N-terminal extension may be partially disordered or only structured in specific functional states, complicating structure-based targeting. Nonetheless, neither region has established precedents for small-molecule inhibition in PGIs. Since, the hook region is situated near the dimer interface and therefore, successfully targeting this motif may disrupt conformational regulation of *Tc* PGI during catalysis thus enabling allosteric modulation with reduced risk of host toxicity.

Given the essential roles of both enolase and PGI in *T. cruzi* metabolism, we performed high-throughput virtual screening of candidate inhibitors using Schrödinger’s SiteMap and PyRx AutoDock. Both enzymes have previously been explored as potential drug targets in a range of pathogenic organisms due to their critical functions in energy metabolism[9, 26]. Although neither target has progressed to clinical trials, multiple studies have demonstrated the feasibility of selectively inhibiting these enzymes [9, 24]. The compounds used in our screens were derived from a previously reported scaffold shown to possess dual inhibitory activity against glycolytic enzymes, including PGI, in *Leishmania mexicana* [32] suggesting that structurally related compounds can target multiple enzymes within a functionally interconnected metabolic pathway. Although *Tc* enolase and *Tc* PGI catalyze distinct reactions, their integration within the glycolytic network raises the possibility that multi-target inhibitors could exert enhanced antiparasitic effects through synergistic disruption of parasite metabolism. Interestingly, our docking results revealed enzyme-specific preferences for distinct compounds (Table 4), likely reflecting differences in active site architecture and binding pocket topology. For *Tc* enolase, the top-ranking inhibitors (Cpd7, Cpd8, and Cpd12) exhibited binding affinities ranging from −7.7 to −8.0 kcal/mol, significantly stronger than that of the substrate 2-PGA (−5.9 kcal/mol). In contrast, *Tc* PGI favored Cpd15, Cpd21, and Cpd14, with docking scores between −7.6 and −7.8 kcal/mol, also outperforming the natural substrate G6P (−6.2 kcal/mol). Notably, prior studies have shown that ST082230, the parent scaffold of these compounds, inhibits *Tc* PGI but lacks selectivity over the human isoform [32]. However, the selectivity profiles of the other top candidates (e.g. Cpd21 and Cpd14) remain uncharacterized. Further biochemical and structural studies will be necessary to determine whether these derivatives can achieve improved selectivity against the parasite enzyme over the human counterpart.

To evaluate the dynamic stability of the top enzyme-ligand complexes, we performed MD simulations (Figure 7). Among all tested systems, the *Tc* enolase-Cpd8 complex exhibited the lowest RMSD value, indicating strong conformational stability over the simulation period. Additionally, RMSF analysis further revealed minimal flexibility in key catalytic residues, suggesting a robust and sustained interaction with the inhibitor (Figure 7). Cpd8 also demonstrated the most favorable binding free energy for Tc enolase (−42.26 kcal/mol), with electrostatic interactions contributing significantly to complex stabilization. This is consistent with prior studies on charged enolase inhibitors in *Mycobacterium tuberculosis* [9]. For *Tc* PGI, Cpd14 and 21 yielded high binding energies suggesting potential viability as novel inhibitors. Collectively, these results reinforce the stability and binding efficacy of the top-ranked compounds and provide a strong foundation for subsequent in vitro validation and structure-guided optimization.

## Conclusion

This study provides a structural and computational characterization of two essential glycolytic enzymes in *T. cruzi* (enolase and PGI) underscoring their potential as druggable targets for Chagas disease. These findings expand upon earlier work in kinetoplastids, including *T. brucei* and *Leishmania* spp., where structural divergence in glycolytic enzymes has been successfully exploited for selective inhibition. Despite this potential, no clinical trials currently target PGI or enolase in parasitic infections, largely due to a historical lack of pathogen-specific structures and concerns over selectivity. Our results lay a foundation for future structure-based drug discovery efforts, including fragment-based screening and optimization of parasite-selective inhibitors.

## Acknowledgements

The SSGCID is funded by Federal funds from the National Institute of Allergy and Infectious Diseases (NIAID), National Institutes of Health (NIH), Department of Health and Human Services, under Contract No.: 75N93022C00036. SSGCID was funded under NIAID Contracts No.: HHSN272201700059C from September 1, 2017 through August 31, 2022; HHSN272201200025C from September 1, 2012 through August 31, 2017; and HHSN272200700057C from September 28, 2007 through September 27, 2012.

