## Supplementary Table 1 for "Structural Basis of Glycolytic Control in *Trypanosoma cruzi*: Insights from Enolase and PGI"

**Supporting information**

**Table S1. *Tc*enolase Macromolecule production information**

| Source organism | *T. cruzi* (strain: CL Brener) |
| --- | --- |
| Forward primer | 5'CTCACCACCACCACCACCATATGACAATCCAAAAGGTTCACGGA3' |
| Reverse primer | 5'ATCCTATCTTACTCACTTACGACCATGCGGGGAAGCCGA3' |
| Expression vector | BG1861 |
| Expression host | *E. coli* BL21 |
| Complete amino acid sequence of the construct produced | MTIQKVHGREILDSRGNPTVEVEVTTELGV FRSAVPSGASTGIHEACELRDDDKRRYLGKG CLNAVKNVNDVLAPALVGK DELQQSTLDKLMRDLDGTPNKSKLGANAIL GCSMAISKAAAARKGVPLYRYLAELAGTKEV RLPVPCFNVINGGKHAGNA LPFQEFMIAPVKAGSFNEALRMGAEVYHSL KSIIKKKYGQDAVNVGDEGGFAPPITDINEP LPILMEAIEQAGHKGRFAI CMDSAASETYDENKKQYNLTFKSPEATWVT AKQLAETYAKWVSEYPIVSLEDPYDQDDFDG FAGITEALKGKAQVVGDDL TVTNVSRIKTAIEKKACNSLLLKINQIGTI TEAIEASKFCMSNGWSVMVSHRSGETEDTYI ADLVVGLGTGQIKTGAPCR GERTAKLNQLLRIEEELGAHAKFGFPAWS |

**Table S2. *Tc* G6PI Macromolecule production information**

| Source organism | *T. cruzi* (strain: CL Brener) |
| --- | --- |
| Forward primer | 5'CTCACCACCACCACCACCATATGAAAGATCAATACCTGAAGGATTT3' |
| Reverse primer | 5'ATCCTATCTTACTCACTTACAAGTGAGAGCGTTCGTTGAATAG3' |
| Expression vector | BG1861 |
| Expression host | *E. coli* BL21 |
| Complete amino acid sequence of the construct produced | MAHHHHHHMKDQYLKDLTVHLNESNAAPAN TSMAVASFNMPHEITRRMRPLGVDADTSLTS CPSWRRLQELYEIHGSESI LKNFDECKDRFQRYSLEVDLRSSDKNFVFL DYSKTHINDEIKDVLFKLVEERGIRAFMRAL FAGEKVNTAENRSVLHIAL RNRSNRPIFVNGHDVMPLVNKVLEQMKKLS EKVRRGEWKGQSGKPIRHVVNIGIGGSDLGP MMACEALRPFSDRRISMHF VSNIDGTHLSEVLNLVDLESTLFIIASKTF TTQETITNALSARNEFLKFLSSRGISEAGAV AKHFVALSTNAEKVKEFGI DEENMFQFWDWVGGRYSLWSAIGLSVMISI GYDNFVELLTGAHIMDEHFINAPTENNLPII LALVGIWYNNFFGSETQAI LPYDQYLWRLPAYLQQLDMESNGKGATKNG RMVSTHTPGIIFGEAGTNGQHAFYQLIHQGT KLIPCDFIGAIQTQNYIGE HHRILMSNFFAQTEALMIGKTPEEVKRELE SAGGKSEDEIQLLIPQKTFTGGRPSNSLLVK ALTPRALGAIIAMYEHKVL VQGAIWGINSYDQWGVELGKLLAKSILLQL QPGQKVTNHDSSTNGLIELFNERSHL |

**Table S3. Crystallization**

|  | ***Tc* enolase** | ***Tc* PGI** |
| --- | --- | --- |
| Method | Vapor diffusion, sitting drop | Vapor diffusion, sitting drop |
| Temperature (K) | 289 | 290 |
| Protein concentration | 47.62mg/ml | 25.0 mg/ml |
| Buffer composition of protein solution | 20 mM HEPES, pH 7.0, 300 mM NaCl, 5% glycerol and 1 mM TCEP | 20 mM HEPES, pH 7.0, 300 mM NaCl, 5% glycerol and 1 mM TCEP |
| Composition of reservoir solution | TrcrA01024aB1 PW35751 CR_27 47.62 mg/ml Morpheus E12 125% *w*/*v* PEG 1000, 125% *w*/*v* PEG 3350, 125% *v*/*v* MPD, 003 *M* of each ethylene glycol 01 *M* bicine/Trizma base pH 85 | Morpheus screen, a8: 12.5% w/v PEG 1000, 12.5% w/v PEG 3350, 12.5% v/v MPD; 30mM of each MgCl2, CaCl2; 0.1 M MOPS/HEPES-Na, TrcrA.17127.a·B1.PS01519 at 25.0 mg ml-1 with 2.5mM glucose-6-phosphate; tray 251762a8, puck rjo4–3; cryo: direct, pH 7.5 |
| Volume and ratio of drop | 0.4 µl, 1:1 | 0.4 µl, 1:1 |
| Volume of reservoir | 80 µl | 80 µl |

**Table S4. Data collection and processing**

|  | ***Tc* enolase** | ***Tc* PGI** |
| --- | --- | --- |
| Diffraction source | ALS beamline 5.0.3 | APS beamline 21-ID-G |
| Wavelength (Å) | 2.4 Å | 1.8 Å |
| Temperature (K) | 100 | 100 |
| Detector | ADSC quantum 315r CCD | RayoniX MX-300 CCD |
| Data collection scaling software | *XSCALE* | *XSCALE* |
| Data reduction software | *XDS* | *XDS* |
| Exposure time per image (s) |  |  |
| Space group | *C*222_1_ | *P21* |
| *a*, *b*, *c* (Å) | 75.29, 119.31, 110.36 | 69.84, 127.03, 72.40 |
| α, β, γ (°) | 90, 90, 90 | 90, 109.74, 90 |
| Mosaicity (°) |  |  |
| Resolution range (Å) | 50.000-2.400 (2.46-2.40) | 50-1.800 (1.850-1.800) |
| Total No. of reflections |  |  |
| No. of unique reflections |  | 109888 |
| Completeness (%) | 99.7 (100.0) | 98.600 (97.900) |
| Redundancy |  | 3.86 |
| 〈 *I*/σ(*I*)〉 | 19.7400 | 13.670 |
| *R*_r.i.m._ |  | 0.089 |
| Overall *B* factor from Wilson plot (Å^2^) | 47.061 | 16.850 |

Values for the outer shell are given in parentheses.

**Table S5.** Ramachandran plot statistics. The statistic values of the Ramachandran plots were generated by the European Bioinformatics Institute’s PDB sum web server.

|  | Number of residues | Percentage (%) |
| --- | --- | --- |
| Most favourable region [A,B,L] | 1000 | 92.0 |
| Additional allowed regions [a,b,l,p] | 87 | 8.0 |
| Generously allowed regions | 0 | 0.0 |
| Disallowed regions | 0 | 0.0 |
| Non-glycine and non-proline residues | 1087 | 100.0 |
| End-residues (excluding Glycine and Proline) | 4 |  |
| Glycine residues | 90 |  |
| Proline residues | 40 |  |
| Total number of residues | 1221 |  |

**Table S6. Docking Scores of Ligands with Enolase (4G7F) and Glucose-6-Isomerase (4QFH)**

| **SN** | **CID** | **NAME** | **Docking Score**  **(kcal/mol)**  ***Tc* enolase** | **Docking Score**  **(kcal/mol)**  ***Tc* PGI** |
| --- | --- | --- | --- | --- |
| 1 | Cpd1 | 2,4-di(aziridin-1-yl)-6-(1-phenyl-1H-pyrrol-2-yl)-1,3,5-triazine | -7.10 | -7.10 |
| 2 | Cpd2 | 2,4-dichloro-6-(1-phenyl-1H-pyrrol-2-yl)-1,3,5-triazine | -6.90 | -6.50 |
| 3 | Cpd3 | 2,4-bis(oxiran-2-ylmethoxy)-6-(1-phenyl-1H-pyrrol-2-yl)-1,3,5-triazine | -7.30 | -7.30 |
| 4 | Cpd4 | 6-(1-phenyl-1H-pyrrol-2-yl)-1,3,5-triazine-2,4-diamine | -7.10 | -7.20 |
| 5 | Cpd5 | N2, N4-dimethyl-6-(1-phenyl-1H-pyrrol-2-yl)-1,3,5-triazine-2,4-diamine | -7.40 | -7.30 |
| 6 | Cpd6 | N2, N2, N4, N4-tetramethyl-6-(1-phenyl-1H-pyrrol-2-yl)-1,3,5-triazine-2,4-diamine | -7.20 | -7.10 |
| 7 | Cpd7 | 2-(1-phenyl-1H-pyrrol-2-yl)-4,6-di(pyrrolidin-1-yl)-1,3,5-triazine | -7.70 | -7.20 |
| 8 | Cpd8 | 2-(1-phenyl-1H-pyrrol-2-yl)-4,6-di(piperidin-1-yl)-1,3,5-triazine | -8.00 | -7.30 |
| 9 | Cpd9 | 1,1'-(6-(1-phenyl-1H-pyrrol-2-yl)-1,3,5-triazine-2,4-diyl) bis(azepane) | -6.90 | -7.10 |
| 10 | Cpd10 | 4,4'-(6-(1-phenyl-1H-pyrrol-2-yl)-1,3,5-triazine-2,4-diyl) dimorpholine | -6.90 | -7.20 |
| 11 | Cpd11 | N, N-dibutyl-4-chloro-6-(1-phenyl-1H-pyrrol-2-yl)-1,3,5-triazin-2-amine | -6.80 | -6.80 |
| 12 | Cpd12 | 4-(azepan-1-yl)-N, N-dimethyl-6-(1-phenyl-1H-pyrrol-2-yl)-1,3,5-triazin-2-amine | -7.90 | -7.40 |
| 13 | Cpd13 | 4-(azepan-1-yl)-N, N-dibutyl-6-(1-phenyl-1H-pyrrol-2-yl)-1,3,5-triazin-2-amine | -7.20 | -7.00 |
| 14 | Cpd14 | 4-(azepan-1-yl)-N, N-dihexyl-6-(1-phenyl-1H-pyrrol-2-yl)-1,3,5-triazin-2-amine | -7.40 | -7.60 |
| 15 | Cpd15 | 4-(azepan-1-yl)-6-(1-benzyl-1H-pyrrol-2-yl)-N, N-dibutyl-1,3,5-triazin-2-amine | -7.00 | -7.50 |
| 16 | Cpd16 | 4-(azepan-1-yl)-N, N-dibutyl-6-(1-phenethyl-1H-pyrrol-2-yl)-1,3,5-triazin-2-amine | -7.10 | -7.40 |
| 17 | Cpd17 | 4-(azepan-1-yl)-N, N-dibutyl-6-(1-(3-phenylpropyl)-1H-pyrrol-2-yl)-1,3,5-triazin-2-amine | -7.40 | -7.10 |
| 18 | Cpd18 | 4-(azepan-1-yl)-N, N-dibutyl-6-(1-(3,5-dimethylphenyl)-1H-pyrrol-2-yl)-1,3,5-triazin-2-amine | -6.50 | -6.90 |
| 19 | Cpd19 | 4-(azepan-1-yl)-N, N-dibutyl-6-(1-cyclohexyl-1H-pyrrol-2-yl)-1,3,5-triazin-2-amine | -7.50 | -7.20 |
| 20 | Cpd20 | 6-(1-benzyl-1H-pyrrol-2-yl)-N2, N2-dibutyl-N4, N4-dihexyl-1,3,5-triazine-2,4-diamine | -6.30 | -6.70 |
| 21 | Cpd21 | 1,1'-(6-(1-(3,5-dimethylphenyl)-1H-pyrrol-2-yl)-1,3,5-triazine-2,4-diyl) bis(azepane) | -6.60 | -7.70 |
| 22 | Substrates | Phosphoglyceric acid/ Glucose-6-Phosphate | -5.90 | -6.20 |

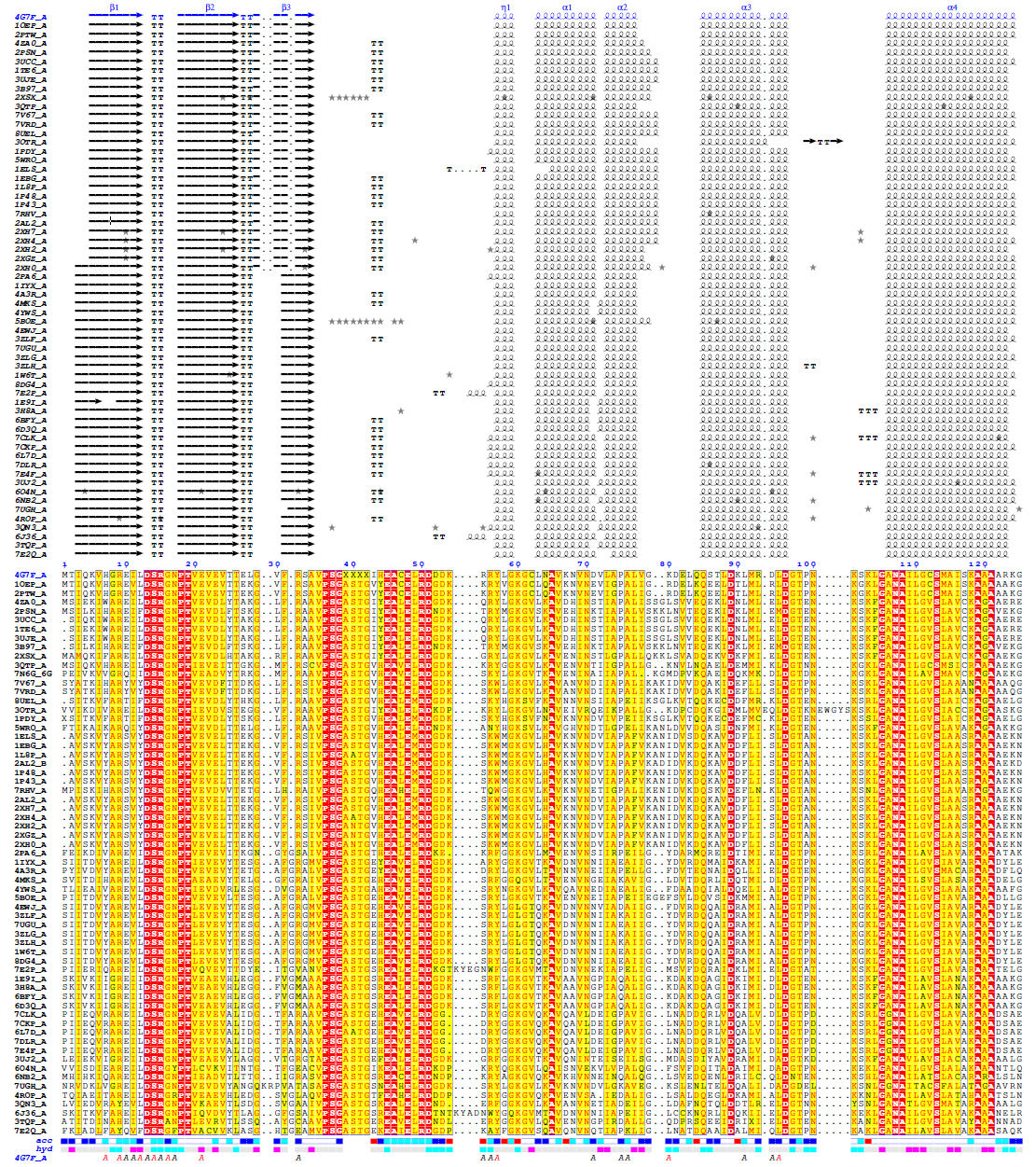

**Fig S1**. *Tc* enolase multiple sequence alignments

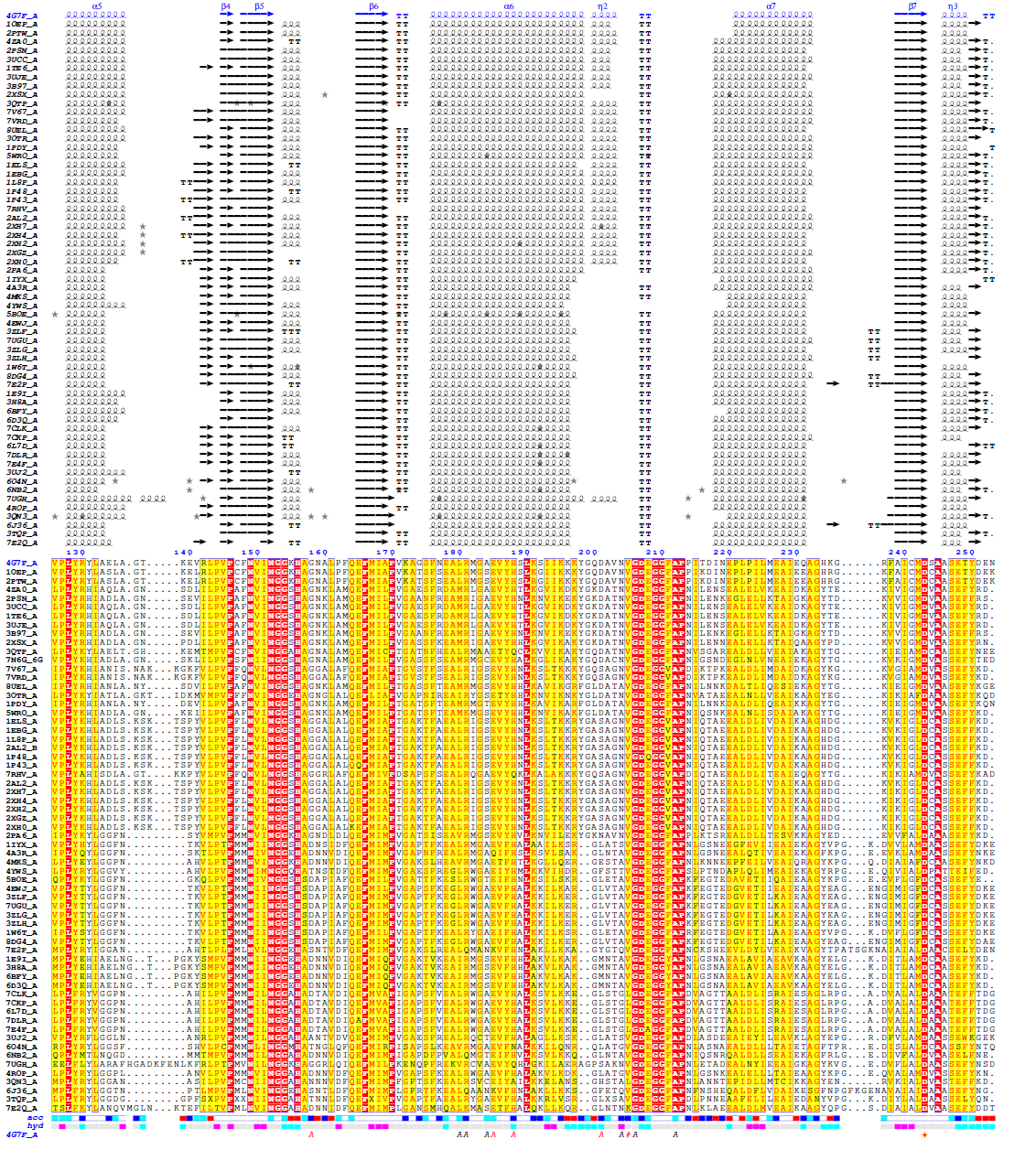

**Fig S2**. *Tc* enolase multiple sequence alignments

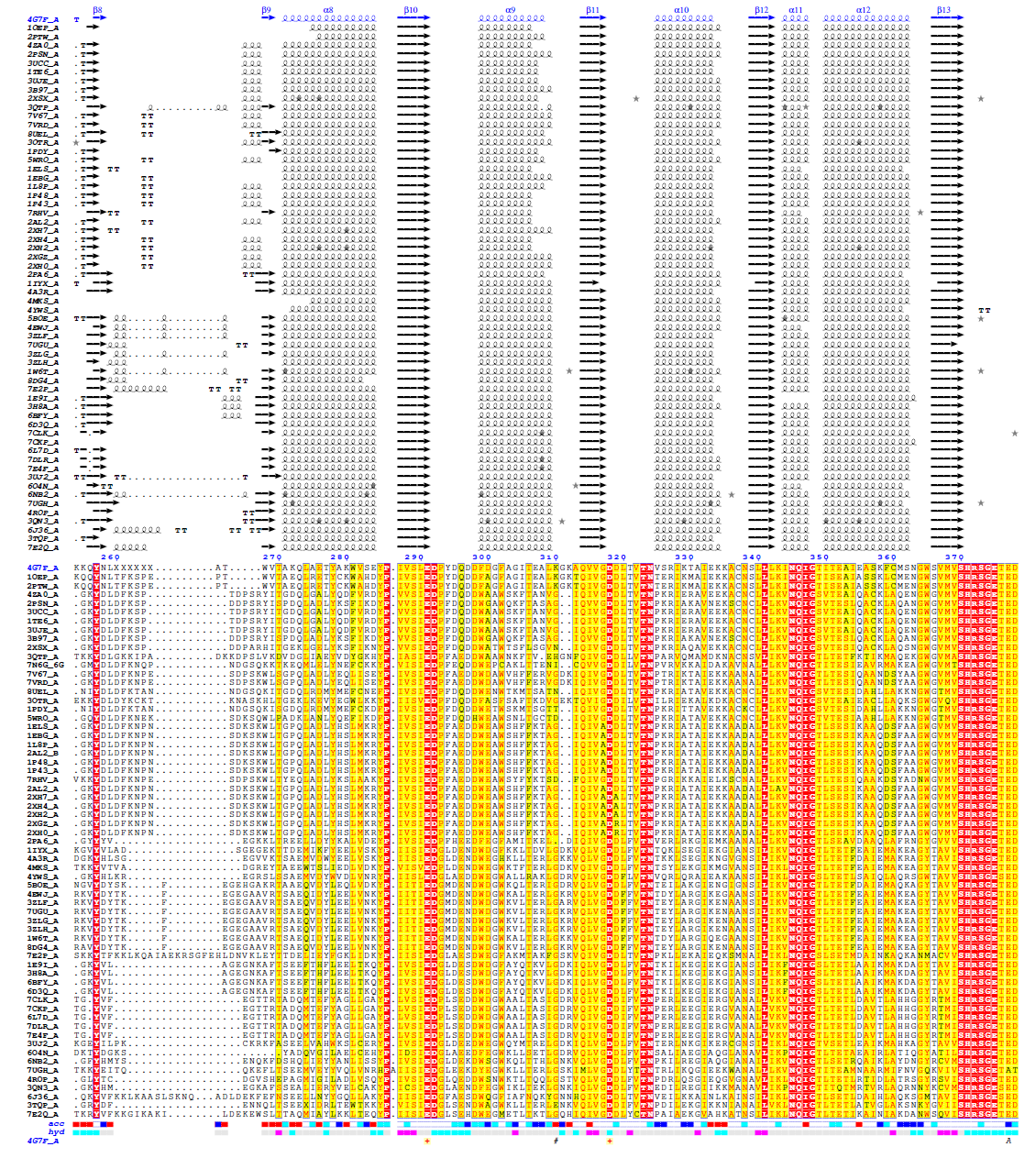

**Fig S3**. Tc enolase multiple sequence alignments

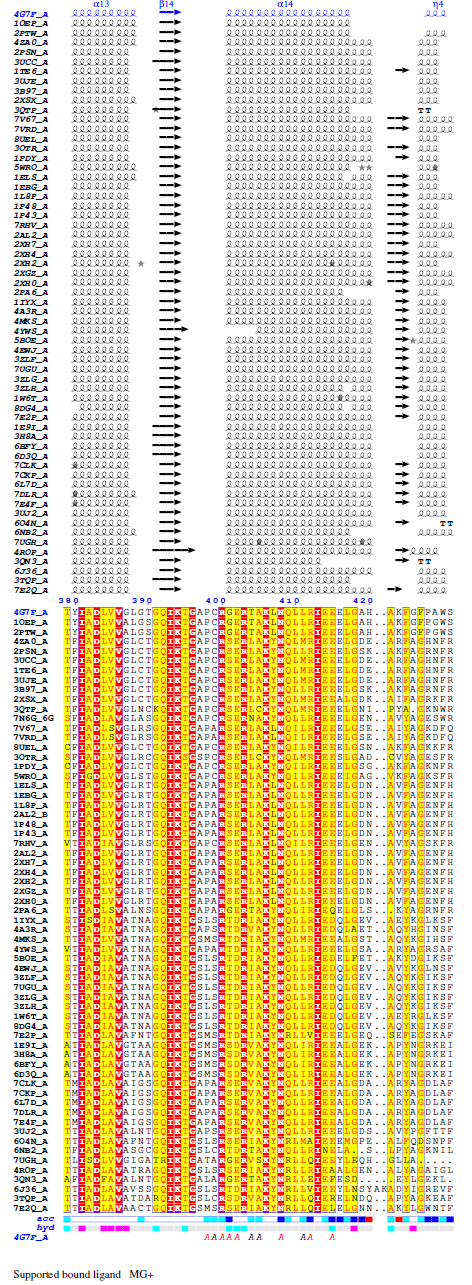

**Fig S4**. Tc enolase multiple sequence alignments

**Fig S5**. Structural comparison of *Tc* enolase (grey) with human enolase 1, 2 and 3.

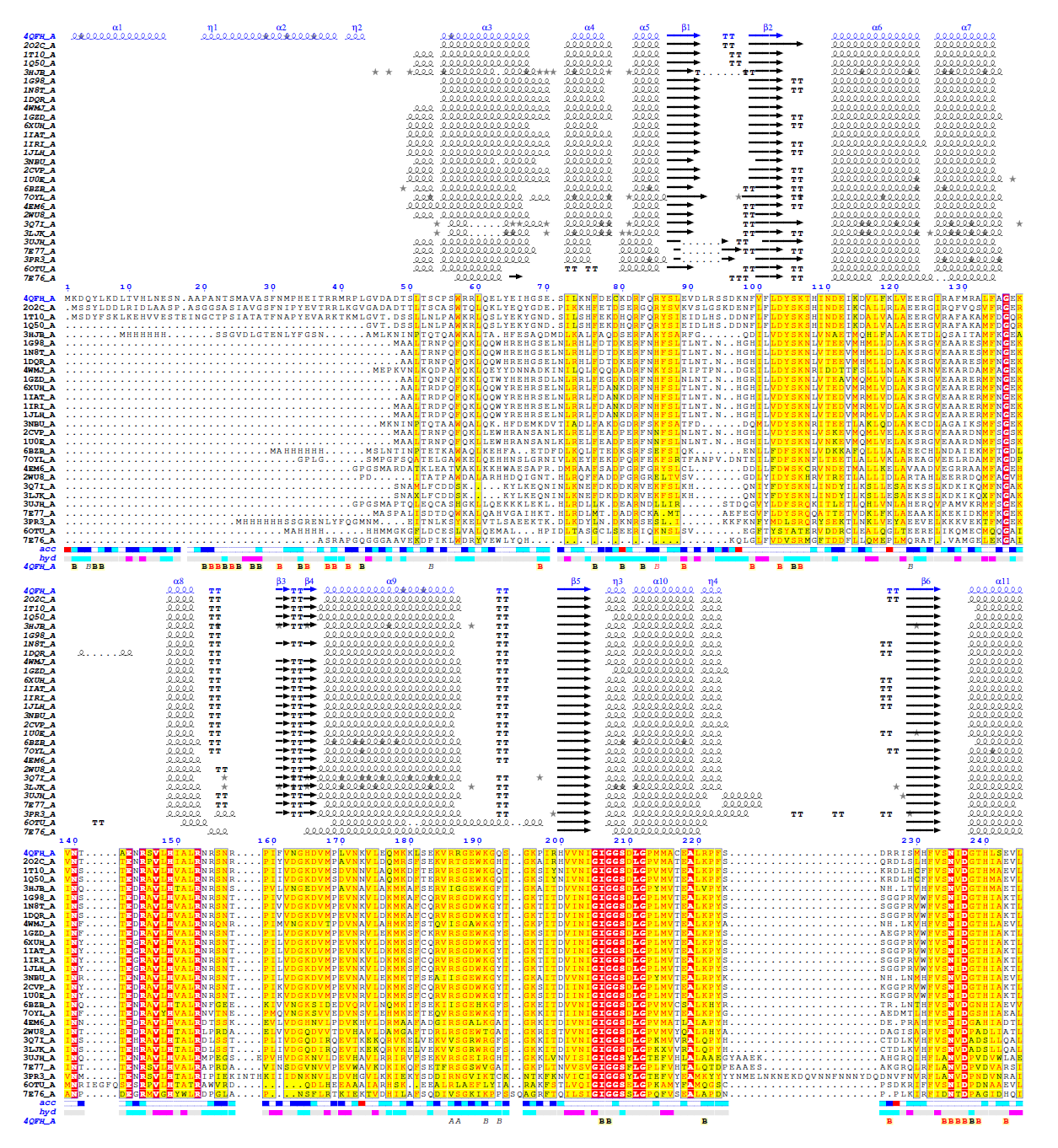

**Fig S6**. *Tc* PGI multiple sequence alignments

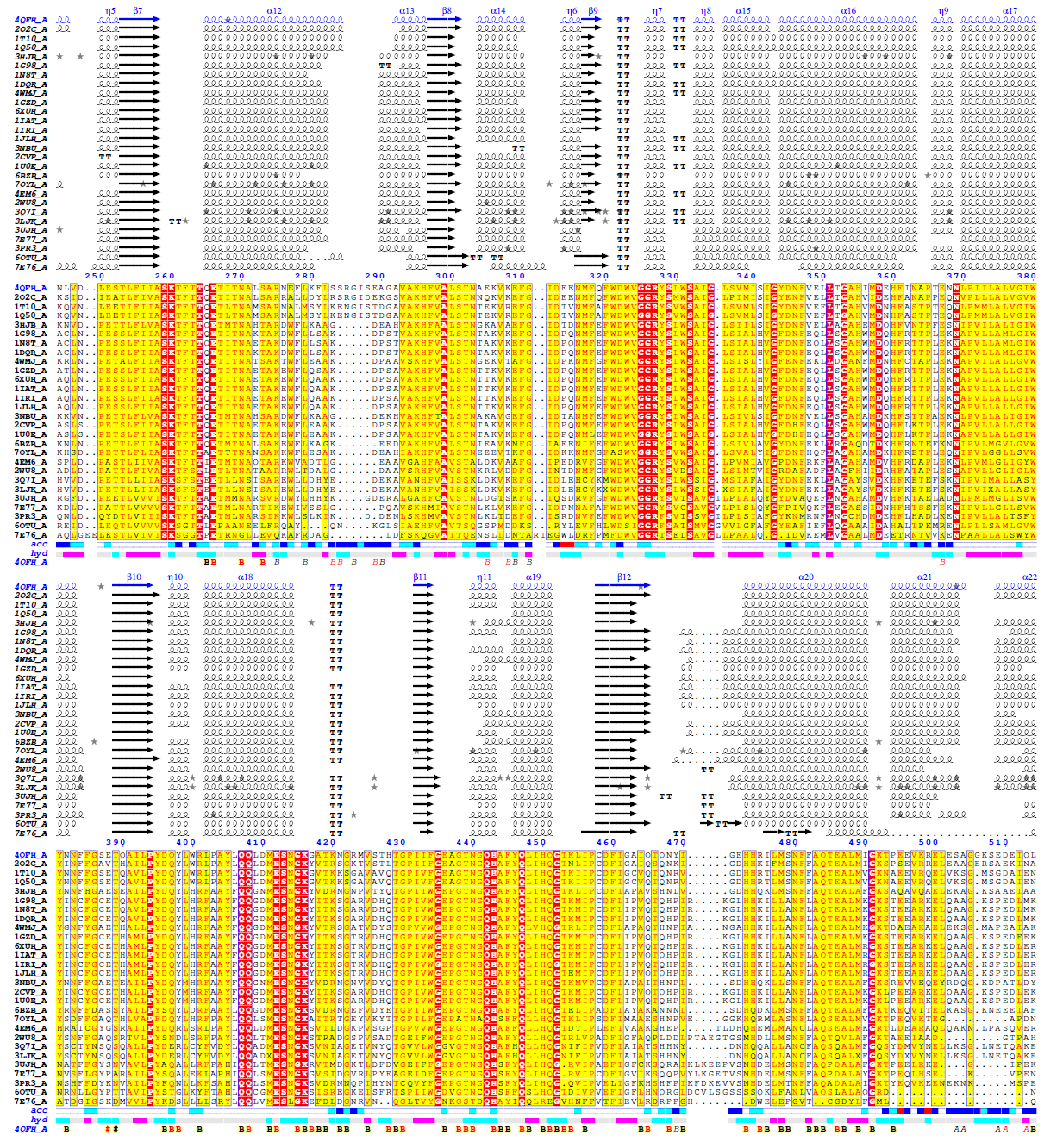

**Fig S7**. *Tc* PGI multiple sequence alignments

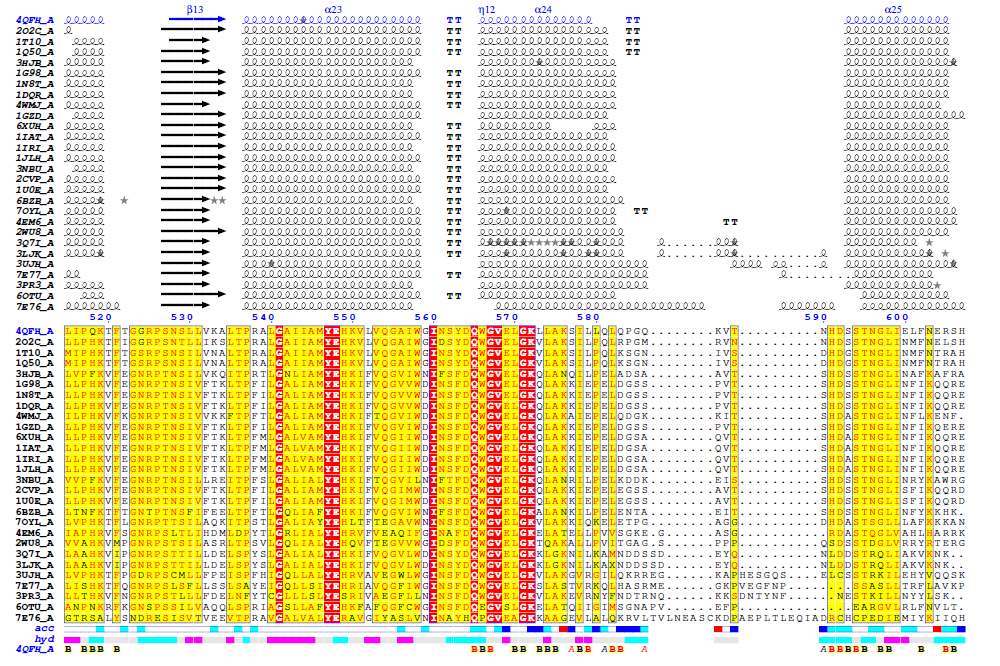

**Fig S8**. *Tc* PGI multiple sequence alignments

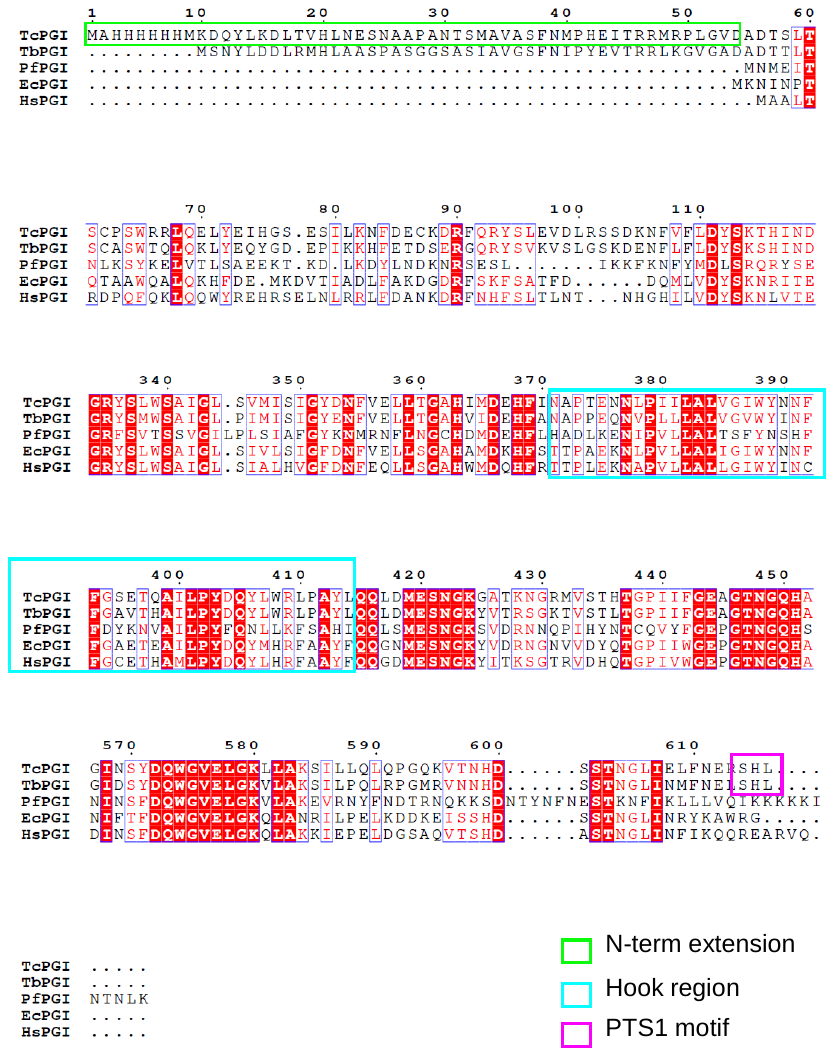

**Fig S9.** Multiple sequence alignments showing unique Tc PGI regions.

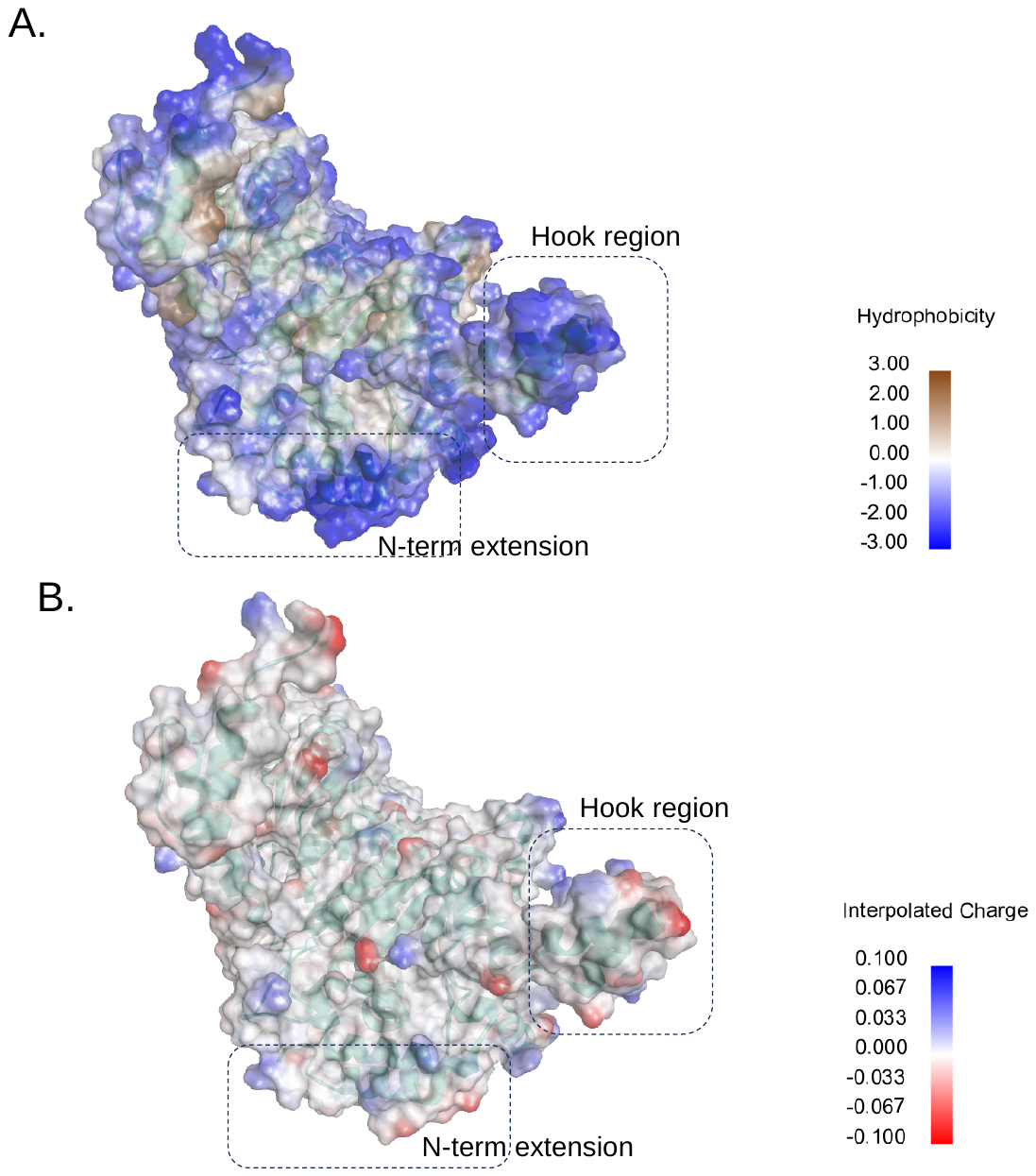

**Figure S10**. Hydrophobic and electrostatic surface properties of Tc PGI. Surface representations colored by hydrophobicity (A) and interpolated electrostatic potential (B). Dashed boxes indicate peripheral regions with distinct physicochemical properties that may influence dimer stability, interfacial interactions, and parasite-specific regulation.

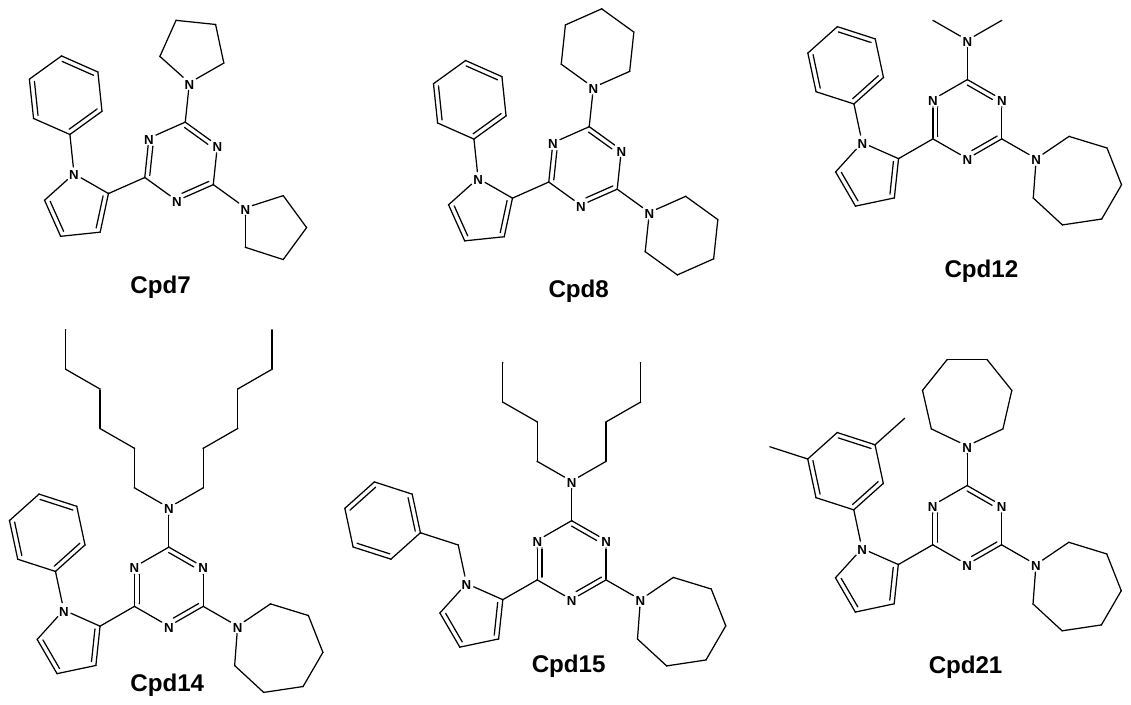

**Figure S11.** Chemical structures of top inhibitors
